# Resolving context-specific protein-protein interactomes for biological discovery and therapeutic target prioritisation

**DOI:** 10.64898/2026.09.08.750135

**Authors:** Alois Thomas, Lisa Fournier, Vincent Jung, Rickie Patani, Pascal Frossard, Raphaëlle Luisier, Cédric Vincent-Cuaz

## Abstract

Protein function is shaped by cellular context, yet most protein representations and interaction maps remain context-agnostic. Here we present ProtScape, a multiscale graph-learning framework integrating global protein interactions, cell-type gene expression and protein language models to learn context-specific representations and infer interactomes across more than 200 cell types. ProtScape substantially outperforms existing approaches in interaction reconstruction, increasing the area under the precision–recall curve by 40 percentage points. Its predicted interactions were supported by held-out continuous STRING global evidence, while its representations recovered higher-order protein organisation. In patient-derived amyotrophic lateral sclerosis motor neurons, ProtScape revealed stage-specific network changes implicating RAB-dependent trafficking as a candidate early disease mechanism. In Parkinson’s disease, it recovered clinically supported therapeutic targets from a proteome-wide search space 16-fold smaller than that required by competing representations. Together, ProtScape provides a scalable framework for translating context-specific interactome organisation into experimentally testable disease mechanisms and therapeutic hypotheses.

## Introduction

Proteins exert their functions through molecular interactions that vary across cell types, physiological states, and disease contexts^1,2,3,4,5,6,7^. Determining how protein-protein interaction (PPI) networks are reorganised across these conditions is therefore essential for understanding protein function and disease-associated molecular changes.

Reference PPI networks catalogue an extensive and still expanding body of experimentally supported interactions, but rarely specify the cellular contexts in which individual interactions occur, resulting in largely context-agnostic interactome maps^3,8,9,10,11,9^. Systematically measuring PPIs throughout the diversity of human cellular environments remains impractical. Filtering a reference interactome using context-specific gene expression is a pragmatic and widely used strategy for approximating cellular PPI networks (Cell-PPIs), across tissues and cell types^12,13,14,15^. In the absence of comprehensive context-resolved proteomic data, this approach leverages widely available transcriptomic measurements as a proxy for protein availability^16^, but necessarily restricts each Cell-PPI to a subgraph of the reference network: it can exclude existing edges, but cannot infer previously unobserved interactions or context-dependent changes in interaction propensity. Cell-type specificity therefore arises from subgraph selection rather than direct inference of context-specific interaction architecture.

Inferring interactions rather than merely filtering them requires machine learning models that couple a protein’s intrinsic molecular properties with its cellular network environment. Protein foundation models (PFMs) capture evolutionary, biochemical and structural constraints from sequence^17,18,19,20^, whereas graph neural networks (GNNs) encode the underlying principles of interaction between proteins to infer new ones from observed network topology^21,22,23,24,25^. Although hybrid PFM–GNN models recently united these complementary signals for PPI prediction, they have largely remained context-agnostic^26,27,28,29,30^. In context-specific settings, Pinnacle recently showed that reference interactions and transcriptomic data can be integrated through GNNs to learn contextual protein representations and interactions across cell types, relying solely on network topology^31,32^.

Here we present ProtScape, a multiscale graph-learning framework spanning protein, cell and tissue organisation that redesigns three core components of context-specific protein modelling to learn contextual representations and infer interactions across more than 200 cell types (Fig. 1). Protscape constructs context-enriched Cell-PPIs that balance network coverage and specificity, preserving higher-order molecular organisation while limiting redundancy across contexts. ProtScape then introduces an asymmetric multiscale architecture that concentrates modelling capacity at the protein level, where context-specific interaction structure is resolved. It combines sequence-derived molecular priors^33^ with context-specific interaction topology through a shared heterophily-aware protein encoder^34^, retaining molecular differences between interacting proteins while identifying interaction-relevant relationships. A masked-interaction learning strategy further requires withheld edges to be reconstructed from protein properties and the surrounding contextual network^35^. To overcome the scarcity of context-resolved functional annotations, we introduce a weakly supervised framework that evaluates contextual protein representations from context-free labels, while identifying the cellular environments that contribute most strongly to each prediction^36,37^.

**Fig. 1.**
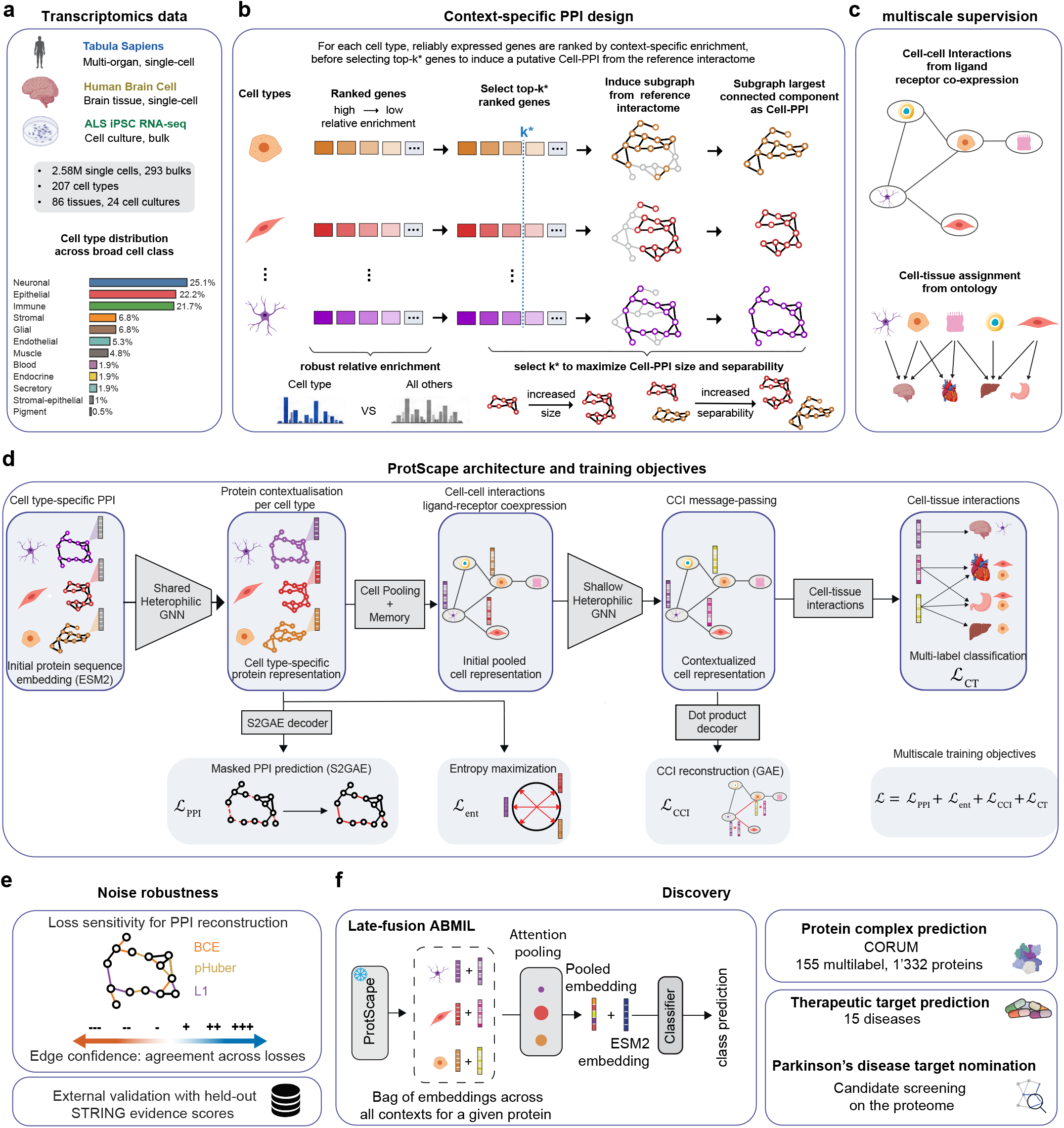
Overview of ProtScape. **a**, Transcriptomic resources used to construct the ProtScape atlas, spanning multi-organ healthy tissues, brain-specific cell populations and ALS-relevant cellular states. **b**, Construction of context-specific protein–protein interaction networks (Cell-PPIs). Reliably expressed genes are ranked by context-specific enrichment, and a context-dependent number of top-ranked genes is selected to derive a connected subgraph of the reference interactome while balancing network size and specificity across cell types. **c**, Multiscale supervision linking cellular contexts through ligand–receptor-supported cell–cell interactions and ontology-derived cell–tissue relationships. **d**, ProtScape architecture and training objectives. Sequence-derived protein representations are contextualised within each Cell-PPI and progressively integrated across cell and tissue scales using shared graph neural networks and multiscale supervision. **e**, Sensitivity of interaction inference to alternative PPI reconstruction objectives, evaluated through changes in predicted edge confidence and comparison with held-out STRING evidence scores. **f**, Weakly supervised evaluation of contextual protein representations for protein-complex prediction and therapeutic-target prioritisation, including proteome-wide candidate nomination in Parkinson’s disease.

ProtScape substantially improves interaction reconstruction over existing approaches and yields context-specific interaction landscapes supported by external biological evidence, together with representations that recover higher-order protein organisation. In patient-derived amyotrophic lateral sclerosis motor neurons, ProtScape identifies disease-stage network changes beyond differential expression and implicates RAB-dependent trafficking as a candidate early disease mechanism. Across therapeutic indications, it improves target prioritisation; in Parkinson’s disease, ProtScape concentrates clinically supported targets within a proteome-wide candidate set that is 16-times smaller than that required by competing models, while nominating new candidates within coherent, disease-relevant molecular systems. Together, these results establish ProtScape as a scalable framework for translating cellular context into experimentally testable disease mechanisms and therapeutic hypotheses.

## Results

### A multiscale framework to learn cell-type-specific protein interactions and representations

Learning contextual protein interactomes requires biologically grounded starting networks. We therefore constructed putative cell-type-specific PPI networks (Cell-PPIs) by restricting a global reference interactome to proteins relevant to each cellular context (Fig. 1a,b). To prioritise these proteins at scale, we used transcriptomic atlases, as cell-resolved proteomics lacks the breadth and depth needed to characterise hundreds of comparable cellular environments. We combined the multi-organ Tabula Sapiens resources^38^ with neuronal populations from the Human Brain Cell Atlas^39^ and iPSC-derived models of amyotrophic lateral sclerosis (ALS)^40^, yielding 207 cellular contexts spanning human tissues, neuronal populations, developmental conditions and disease-associated states, organised into 12 broad cellular categories (Fig. 1a). For each context, we retained reliably expressed genes and ranked them by enrichment relative to other cellular environments^12,41,42^. Each Cell-PPI was then constructed as a subgraph of the reference interactome containing the top-ranked proteins, with their number selected to balance context specificity, network size and connectivity (Fig. 1b; Methods). The resulting Cell-PPIs were large sparse networks (5473 proteins with 37916 interactions on average) that retained broad proteome coverage while remaining distinct across contexts (Extended Data Fig. 1a–d). We next coupled the otherwise independent Cell-PPIs through a metagraph that connected cell types by statistically supported ligand–receptor relationships^43^ and linked them to tissues through ontology-based mappings^44^ (Fig. 1c; Extended Data Fig.1e,f). This established a multiscale protein–cell–tissue scaffold incorporating intercellular communication and anatomical organisation.

ProtScape used this scaffold to jointly learn context-specific protein interactions and representations through an asymmetric multiscale architecture that concentrates modelling capacity at the protein level (Fig. 1d; Methods). Each protein was initialised with an ESM2 representation^33^, providing an intrinsic molecular prior that was subsequently contextualised within each Cell-PPI. Relative to these sequence-derived features, the Cell-PPIs were strongly heterophilic (mean node heterophily: 84.9%^45^), indicating that interacting proteins frequently differed in their intrinsic molecular profiles. ProtScape therefore employed a heterophily-aware Adaptive Channel Mixing (ACM) encoder to preserve protein-specific molecular information while learning interaction-relevant relationships^34^, rather than a conventional message-passing architecture designed primarily for homophilic graphs^46^. Instead of fitting separate GNNs to individual contexts^31,32^, ProtScape jointly optimised a single protein encoder across all Cell-PPIs, enabling information transfer across biological environments while retaining their specificity through the distinct protein composition and topology that compose them. Protein representations within each Cell-PPI were pooled to generate cell-type representations. A single-layer ACM encoder then refined these into contextual cell representations capturing cell–cell interactions and cell–tissue assignments.

ProtScape further replaced conventional full-graph autoencoding^47^, which exposes the encoder to all training edges, with S2GAE-based self-supervised masked edge learning^35^. During training, a subset of PPI edges was removed before encoding and reconstructed using only the retained topology and molecular features. The model was therefore required to infer missing interactions rather than reproduce observed network structure. This reconstruction objective was combined with regularisation of contextual protein representations and higher-order cell-cell and cell-tissue supervision. Together, these objectives decoupled context definition from interaction inference: transcriptomic enrichment specified the initial Cell-PPI scaffolds, whereas molecular priors, masked edge learning and metagraph supervision drove the inference of context-specific interactions and the learning of contextual protein and cell representations. We next assess the robustness and biological support of these inferred interaction landscapes (Fig. 1e), before determining whether the resulting representations capture higher-order protein organisation and reveal disease-associated mechanisms and therapeutic opportunities (Fig. 1f).

### ProtScape predicts biologically supported interactions across cell types

To determine whether ProtScape learned generalisable interaction structure rather than memorising its input networks, we first assessed its recovery of held-out Cell-PPI edges and then examined whether newly predicted interactions received external biological support.

We compared ProtScape with Pinnacle retrained on the same Cell-PPIs^31,32^, Pinnacle variants incorporating ESM2 features^33^ or heterophily-aware message passing^34^, and ProtScape-GAE, in which S2GAE-based masked edge learning was replaced by conventional graph autoencoding^47^ (Fig. 2a; Methods). To reflect the sparsity of biological interaction networks, models were evaluated at increasing negative-to-positive edge ratios. Finally, each unique protein pair was assigned exclusively to either the training or testing set to prevent cross-context leakage (Methods).

**Fig. 2.**
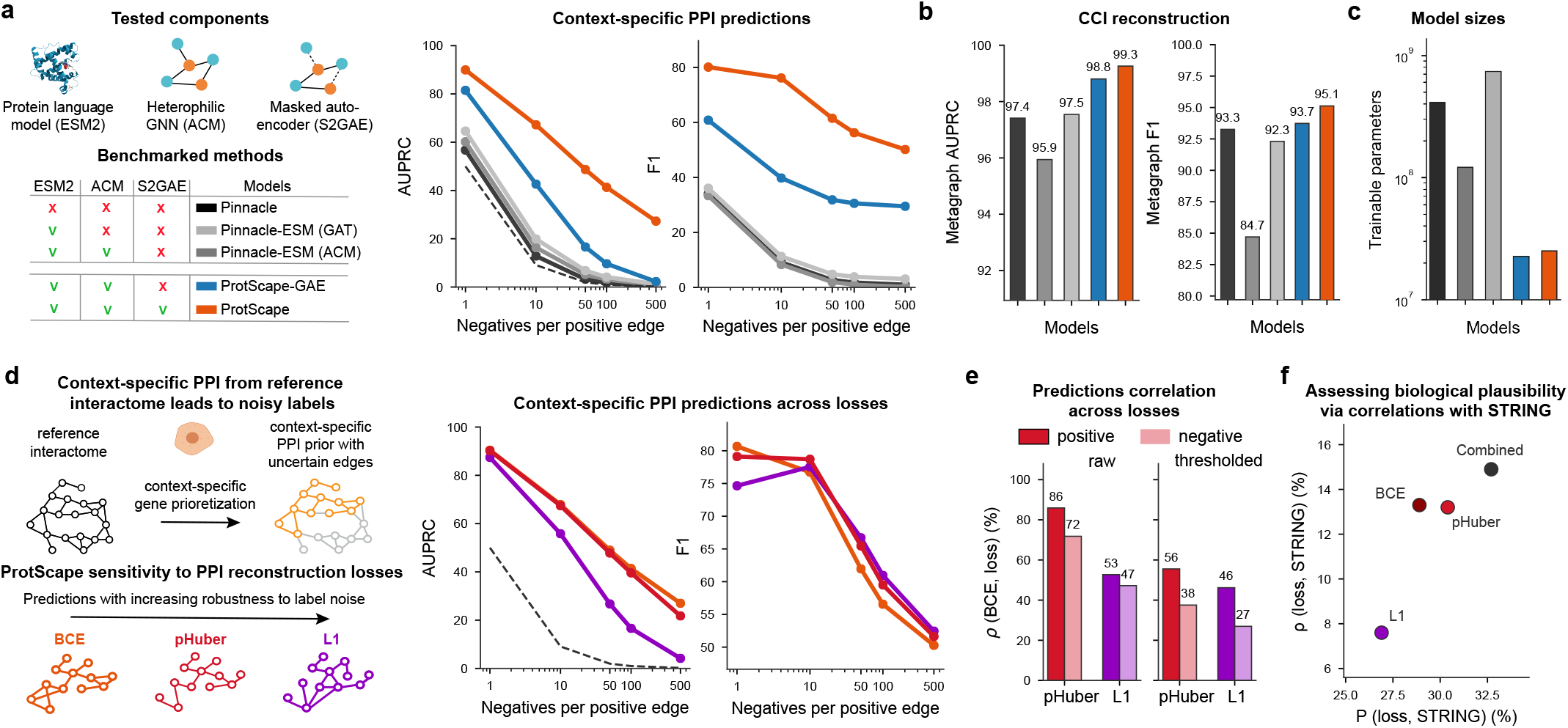
ProtScape predicts biologically supported context-specific protein interactions under putative Cell-PPI supervision. **a**, Held-out context-specific PPI prediction across increasing negative-to-positive edge ratios. We compare Pinnacle and variants incorporating ESM2 representations or heterophily-aware Adaptive Channel Mixing (ACM) with ProtScape-GAE and full ProtScape; the component matrix indicates the methodological elements included in each model. Performance is reported as AUPRC and macro-F1; dashed lines indicate AUPRC from random classifiers. **b**, Held-out reconstruction of cell–cell interactions in the metagraph, evaluated by AUPRC and macro-F1. **c**, Number of trainable parameters for each model, shown on a logarithmic scale. **d**, Sensitivity of ProtScape interaction prediction to the PPI reconstruction objective. Cell-PPIs derived from context-specific gene prioritisation provide putative interaction labels; otherwise, we trained identical models using BCE, pHuber, or L1 losses, which differ in sensitivity to prediction–label discrepancies. We evaluate held-out interaction reconstruction across increasing class imbalance using AUPRC and macro-F1. **e**, Agreement between predictions obtained with alternative reconstruction objectives. Spearman (*ρ*) correlations between BCE and pHuber or L1 predictions are shown for positive and negative Cell-PPI labels before and after thresholding. **f**, External assessment of the inferred interaction landscapes using held-out continuous STRING confidence scores. Pearson (P) and Spearman (*ρ*) correlations are reported for predictions obtained with BCE, pHuber, L1 and their combination, capturing complementary aspects of agreement with STRING evidence.

ProtScape consistently achieved the highest AUPRC^48^ and macro-F1^49^ across cellular contexts and increasingly stringent class-imbalance settings (Fig. 2a; Extended Data Fig. 2a). ProtScape-GAE alone surpassed all Pinnacle variants, indicating that the multiscale architecture better exploits the context-specific interactome topology, independent of masked edge learning. Incorporating the latter further increased performance, with the advantage of full ProtScape increasing as the candidate interaction space became more imbalanced. These gains extended across biological scales, with ProtScape more accurately reconstructing cell-cell interactions despite a comparatively simple metagraph architecture (Fig. 2b). Remarkably, these results were achieved with about 20 times fewer trainable parameters than Pinnacle-based models (Fig. 2c).

Because the initial Cell-PPIs were computationally constructed rather than experimentally measured, improved reconstruction could reflect closer reproduction of their binary structure rather than greater biological plausibility. Likewise, interactions absent from the initial Cell-PPIs could represent either false positives or plausible interactions missing from the starting networks. We therefore assessed biological plausibility in two steps. First, we trained otherwise identical ProtScape models using BCE, pHuber and L1 reconstruction objectives, which progressively reduce the penalty for large prediction–label discrepancies, and thereby vary the model’s dependence on the initial networks (Fig. 2d; Methods). BCE and pHuber achieved nearly identical reconstruction performance, whereas L1 showed decreasing AUPRC as class imbalance increased despite comparable macro-F1. Similar aggregate performance, however, did not imply identical predictions, with BCE and pHuber showing the greatest concordance and L1 the least (Fig. 2e; Extended Data Fig. 2d). We then evaluated the biological plausibility of these landscapes by comparing predictions from each reconstruction objective with continuous STRING association scores, cumulating experimental, computational and literature evidence^50^. Approximately 60% of the reference interactions were represented in STRING (Extended Data Fig. 2h); however, ProtScape was trained only on binary Cell-PPI labels and had no access to the continuous STRING scores. Because STRING does not resolve associations by cellular context, we summarised each protein pair by the mean and variability of its ProtScape scores across cell types and related these quantities to STRING evidence (Methods). Across reconstruction objectives, STRING scores increased with mean prediction strength but decreased with contextual variability (Fig. 2f), indicating stronger global support for interactions predicted broadly across cellular environments than for context-restricted predictions. This pattern is consistent with STRING integrating evidence across studies and biological settings, such that recurrent interactions are more likely to accumulate high scores. BCE and pHuber showed the strongest agreement with STRING, indicating that the objectives that most effectively reconstructed the initial Cell-PPIs also produced interaction landscapes most consistent with external graded evidence (Fig. 2f; Extended Data Fig. 2c-i).

Together, these results establish ProtScape as a robust framework for learning biologically supported context-specific interaction landscapes from putative Cell-PPI scaffolds without being constrained to reproduce their binary topology.

### Contextual protein representations capture higher-order complex organisation

Many cellular functions rely on proteins acting cooperatively within molecular complexes. We next tested whether ProtScape representations capture higher-order organisation beyond pairwise PPIs by predicting membership in experimentally characterised CORUM^51^ complexes (Extended Data Fig. 3a, Methods). ProtScape represents each protein across multiple Cell-PPIs, whereas CORUM provides protein-level annotations without specifying cellular context. We therefore developed an attention-based multiple-instance learning (ABMIL^36^) framework that integrates the full set of contextual representations and identifies the contexts that contribute to each prediction (Fig. 3a). Freezing the pretrained encoders ensured that performance reflected information acquired during pretraining rather than adaptation to CORUM labels.

**Fig. 3.**
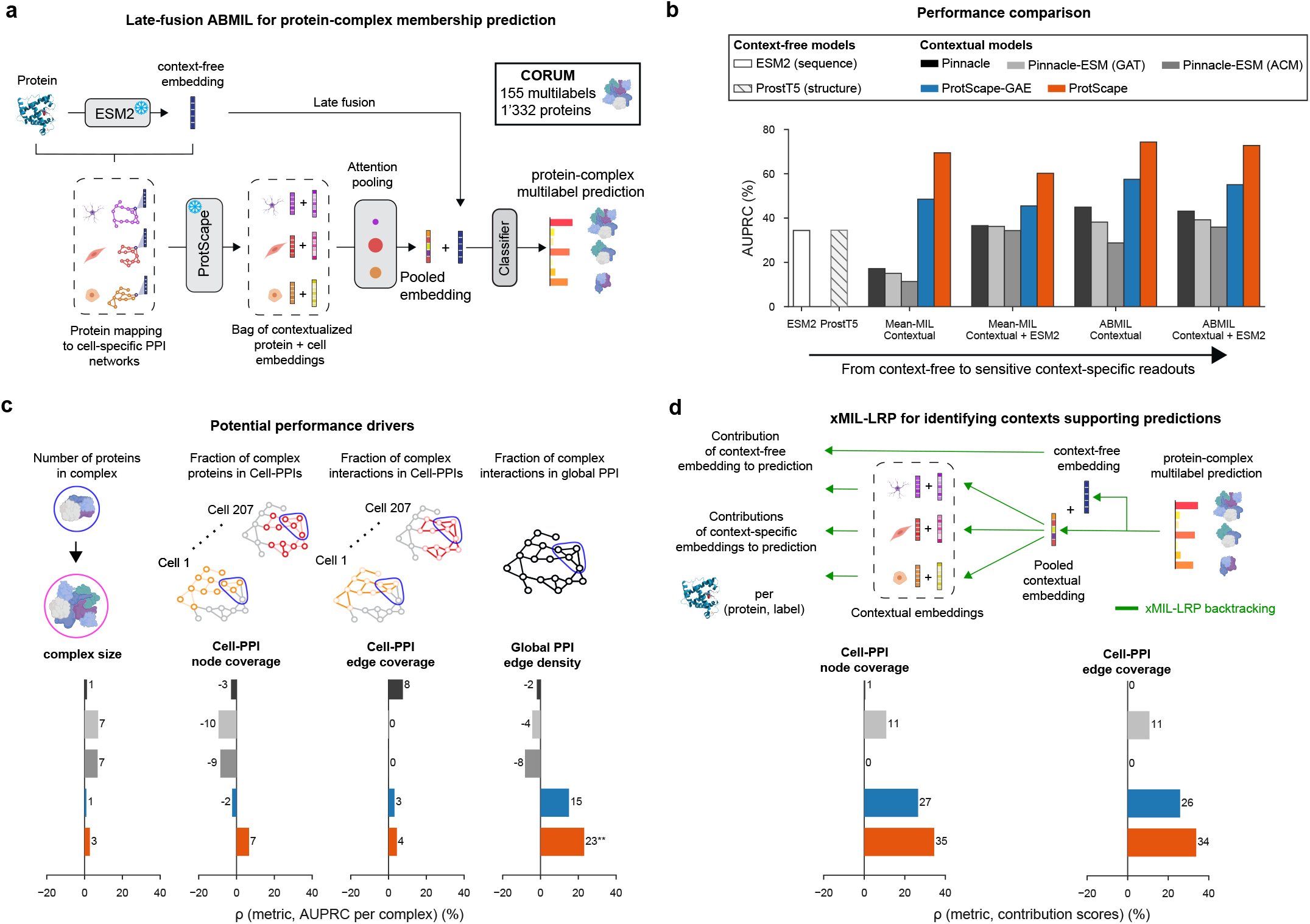
ProtScape captures higher-order protein organisation and identifies cellular contexts supporting protein-complex predictions. **a**, Weakly supervised evaluation of 155 human protein complexes from CORUM ^51^. Each protein is represented by its embeddings across cellular contexts, which are aggregated by attention-based multiple-instance learning (ABMIL) and optionally combined with a context-free ESM2 representation before multilabel complex prediction. Protein encoders remain frozen during downstream evaluation. **b**, Protein-complex prediction performance for context-free ESM2 and ProstT5 representations and contextual Pinnacle, ProtScape-GAE and ProtScape representations across alternative context-aggregation and ESM2-fusion strategies. Bars show test AUPRC macro-averaged across the 155 complexes and averaged across five models selected by five-fold cross-validation on the non-test proteins. **c**, Associations between protein-complex prediction performance and four predefined structural properties: complex size, mean complex-protein coverage across the 207 Cell-PPIs, mean within-complex edge coverage across Cell-PPIs and within-complex edge density in the global reference interactome. Bars report Spearman correlations between each metric and per-complex performance. **(P<0.01); two-sided Spearman tests, unadjusted. **d**, Context-level interpretation of protein-complex predictions using xMIL-LRP. Predictive relevance was backpropagated to individual contextual protein representations, and the resulting contribution scores were related to the representations of the corresponding complex proteins and interactions within each Cell-PPI. Bars report median within-complex Spearman correlations across eligible complexes.

ProtScape achieved the highest protein-complex prediction performance, and ABMIL consistently outperformed simpler context-aggregation strategies (Fig. 3b; Extended Data Fig. 3b). ProtScape-GAE surpassed all Pinnacle variants, while masked S2GAE pretraining further improved functional transfer. Simply combining ESM2 embeddings with averaged ProtScape representations did not improve performance, whereas ABMIL effectively integrated sequence and contextual information by selectively aggregating the cellular environments relevant to each protein. Performance also depended on the reconstruction objective used during pretraining, with BCE yielding the strongest results (Extended Data Fig. 3c).

We next examined how the ProtScape–ABMIL framework combines contextual protein representations to recover protein-level complex annotations. We thus related per-complex performance to complex size, coverage within individual Cell-PPIs and interaction coverage in the global reference interactome (Fig. 3c). Only reference-interactome edge density was significantly associated with performance, suggesting that prediction drew on interaction structure distributed across the Cell-PPI atlas rather than requiring a complex to be concentrated within a single context. Nevertheless, context-specific contributions obtained from xMIL-LRP^37^ revealed greater positive contributions from Cell-PPIs retaining larger fractions of the corresponding complex proteins and interactions (Fig. 3d; Methods). These contexts represent plausible cellular environments for complex assembly, as they are associated with reliably expressed proteins that are specifically enriched within these environments.

Together, these results show that ProtScape representations encode higher-order protein organisation distributed across cellular contexts. ABMIL recovers and localises this information using only protein-level annotations, providing a general strategy for evaluating and interpreting contextual protein representations without context-resolved supervision.

### ProtScape nominates ALS-associated network rewiring linked to synaptic dysfunction

Protein-interaction networks can change in disease without corresponding changes in transcript or protein abundance, potentially revealing mechanisms that differential-expression analysis misses. The ProtScape training atlas included Cell-PPIs derived from control and ALS-associated VCP-mutant patient-derived iPSCs sampled throughout motor-neuron differentiation^52,53,54^ (Fig. 4a,b). We therefore asked whether the learned interaction landscapes revealed network changes as motor neurons underwent terminal differentiation and VCP-mutant cultures began to manifest disease-associated phenotypes, thereby focusing on days 22 and 35.

**Fig. 4.**
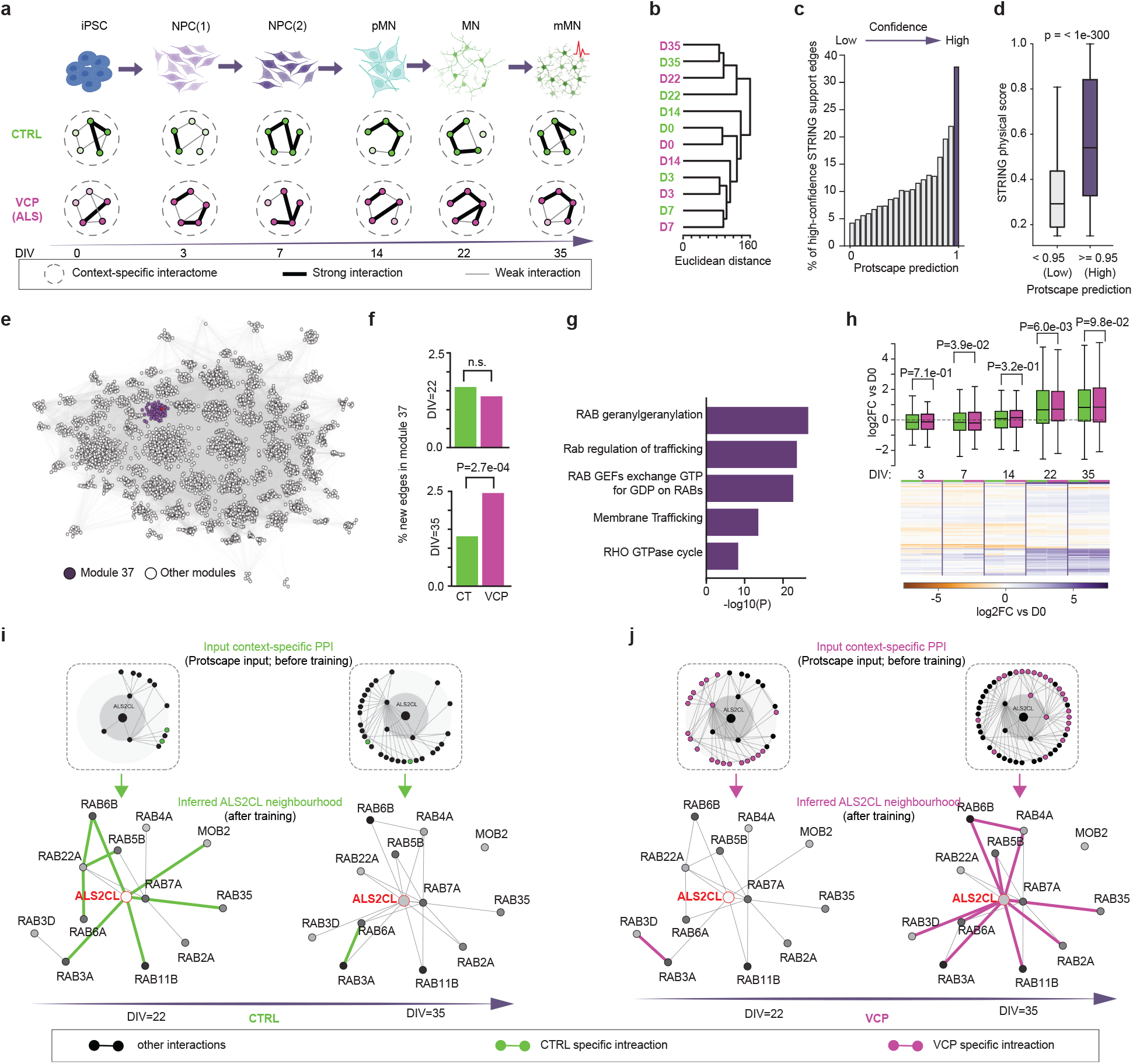
Context-specific interactome modelling reveals ALS-associated network rewiring during motor-neuron differentiation. **a**, Longitudinal differentiation of control (CTRL) and VCP-mutant ALS patient-derived iPSCs through neural progenitor (NPC), precursor motor-neuron (pMN), motor-neuron (MN) and mature motor-neuron (mMN) states ^40^. Context-specific protein interaction networks (Cell-PPIs) were constructed at each differentiation stage. Edge width illustrates interaction strength. **b**, Similarity of inferred cell-PPIs across differentiation stages and conditions, measured by hierarchical clustering of ProtScape predictions, using Euclidean distance and complete linkage. **c**, Percentage of novel interactions with high-confidence STRING support (score >0.75) within successive 0.05-wide bins of ProtScape scores. Novel interactions were defined as protein pairs absent from the global reference interactome; only pairs with a non-zero STRING physical-interaction score were included. **d**, STRING physical-interaction scores for novel interactions with ProtScape scores below or above the threshold of 0.95 used in subsequent analyses. Centre lines indicate medians, boxes indicate interquartile ranges and whiskers extend to 1.5 times the interquartile range. The indicated P value was calculated using a two-sided Mann–Whitney U test. **e**, Network comprising genotype-selective novel interactions with ProtScape scores 0.95 and reference interactions connecting proteins incident to these edges. The network was partitioned into 80 communities using iterative Leiden clustering; module 37 is highlighted. **f**, Percentage of all CTRL- or VCP-selective novel interactions assigned to module 37 at DIV 22 and 35. Genotype-specific enrichment was evaluated using two-sided Fisher’s exact tests, with Benjamini–Hochberg correction performed separately at each time point; n.s., not significant. **g**, Reactome terms enriched among proteins in module 37. Bars show log10-transformed FDR-adjusted enrichment P-values. **h**, Gene expression dynamics of proteins belonging to module 37 across CTRL and VCP-mutant motor-neuron differentiation. Expression is shown as log2 fold change relative to the corresponding genotype-specific DIV-0 value. Centre lines indicate medians, boxes indicate interquartile ranges, and whiskers extend to 1.5 times the interquartile range; each observation represents one module gene. CTRL and VCP values were compared at each time point using two-sided paired Wilcoxon signed-rank tests. The heat map shows gene-level expression changes across differentiation. **i**,**j**, Context-specific network topology and inferred ALS2CL interaction neighbourhoods in module 37 in CTRL (i) and VCP-mutant (j) cells at days 22 and 35. Insets show the initial context-specific PPI neighbourhoods used as input to ProtScape before training; lower networks show the ALS2CL-centered neighbourhood in module 37 after context-specific interaction inference. Grey edges represent other interactions, whereas green and magenta edges denote high-confidence interactions selectively predicted in CTRL and VCP-mutant cells at the indicated developmental stage, respectively.

To identify high-confidence interactions absent from the input Cell-PPIs, we calibrated ProtScape edge scores against STRING physical-interaction evidence. The proportion of edges with high-confidence STRING support (physical score >0.75) increased progressively with ProtScape score, with a pronounced rise among the highest-scoring predictions (>0.95). We therefore used this threshold to identify condition-selective candidate novel interactions in control and VCP-mutant motor neurons (Fig. 4c,d). Combining these predictions with reference interactions produced a condition-resolved network comprising 80 functionally coherent modules (Fig. 4e; Extended Data Fig. 4a), several of which showed stage- and genotype-selective remodelling (Extended Data Fig. 4b). Module 37 was particularly enriched for VCP-selective predictions at day 35 and converged on RAB-dependent membrane trafficking (Fig. 4f,g).

Within module 37, ALS2CL provided a potential link to endosomal trafficking, a pathway broadly implicated in ALS and other neurodegenerative diseases^55,56^. ALS2CL binds RAB5 and regulates endosomal dynamics, and it interacts with ALS2/alsin, a RAB5 guanine-nucleotide exchange factor mutated in recessive motor-neuron disorders^57,58^.

Although ALS2-associated disease differs from dominantly inherited VCP-ALS in its genetic basis, clinical presentation and the motor-neuron populations primarily affected, the predicted ALS2CL neighbourhood contained multiple RAB-family proteins and underwent pronounced genotype-dependent remodelling between days 22 and 35 (Fig. 4i,j), suggesting that these distinct forms of ALS converge on endosomal trafficking.

This pattern was not explained by differential expression alone. Module proteins were broadly retained across conditions and showed comparable expression between control and VCP-mutant cells at each stage (Fig. 4h; Extended Data Fig. 4d), yet the local topology and predicted one- and two-hop neighbourhoods of ALS2CL differed by stage and genotype (Fig. 4i,j; Extended Data Fig. 4c).

ProtScape therefore extended contextual differences in the input networks to nominate additional disease-associated relationships absent from the initial Cell-PPIs. Together, these analyses nominate stage-specific remodelling of an ALS2CL–RAB trafficking programme as a testable mechanism contributing to early synaptic dysfunction in VCP-mutant motor neurons.

### Context-specific protein representations improve therapeutic-target prioritisation across diseases

Therapeutic targets must act within disease-relevant cellular environments, yet their effects may emerge from molecular processes distributed across multiple cell types and tissues^59^. We therefore assessed whether ProtScape could improve disease-specific target prioritisation, identify the contexts supporting individual predictions, and narrow the proteome-wide search for candidate targets.

Extending Pinnacle’s evaluation from 2 to 15 diseases across five therapeutic areas, models distinguished targets supported by indication-specific drugs that had completed at least phase 2 trials from druggable proteins lacking comparable therapeutic evidence (Fig. 5a; Methods). Because these annotations were defined at the protein level, we used ABMIL to integrate each protein’s contextual representations and identify the cell types contributing to its prediction. ProtScape-ABMIL achieved the strongest overall performance, with particularly strong performance in Parkinson’s disease (PD), indicating that ProtScape provides complementary disease-relevant information beyond sequence-only representations (Fig. 5b,c; Extended Data Fig.5a,b). Context attributions were also biologically coherent: immune-cell representations contributed most strongly to inflammatory and autoimmune indications, whereas neuronal, glial as well as muscle and stromal contexts predominated in PD (Extended Data Fig. 5c).

**Fig. 5.**
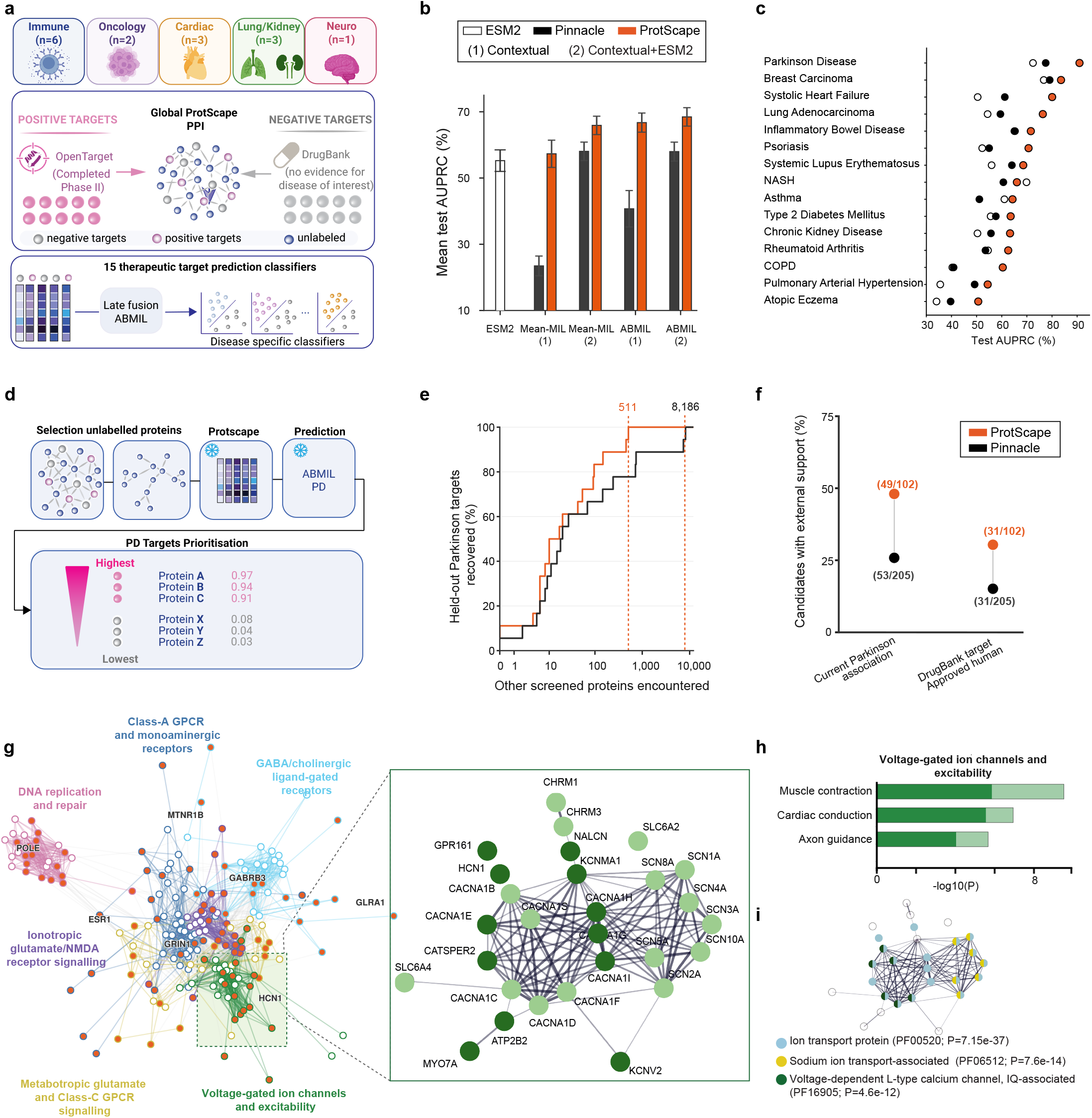
ProtScape improves therapeutic-target prioritisation and identifies biologically coherent Parkinson’s disease candidates. **a**, Therapeutic-target prediction across 15 diseases spanning five therapeutic areas. Open Targets-supported targets (phase 3 or later clinical evidence, or phase 2 evidence with completed status) are contrasted with druggable proteins lacking indication-specific evidence; disease classifiers use frozen ProtScape representations with late-fusion ABMIL. **b**, Mean test AUPRC across diseases for ESM2 and contextual models using alternative aggregation and ESM2-fusion strategies; error bars show s.e.m. **c**, Disease-specific test AUPRC for ESM2, Pinnacle and ProtScape. **d**, Proteome-wide Parkinson’s disease (PD) prioritisation, applying the trained PD classifier to global-interactome proteins to rank previously unlabelled candidates. **e**, Screening depth required to recover 18 held-out clinically supported PD targets. Curves show cumulative recovery; dashed lines mark complete recovery by ProtScape (511 proteins) and Pinnacle (8,186 proteins). **f**, External support for ProtScape- and Pinnacle-nominated PD candidates. Points show the fraction with current Open Targets PD associations or approved human DrugBank-target annotations; labels indicate supported versus total nominated candidates. **g**, STRING network of ProtScape-nominated candidates and established PD targets, revealing disease-relevant functional modules; the inset highlights a voltage-gated ion-channel and neuronal-excitability module. **h**, Functional enrichment of this module, highlighting cardiac conduction, muscle contraction and axon guidance. **i**, Pfam-domain organisation within the module; colours denote enriched ion-transport and voltage-gated ion-channel domains.

We next asked how far into the proteome each model had to search to recover the 18 clinically supported PD targets held out during training. ProtScape recovered all 18 within the top approximately 500 proteins, whereas Pinnacle required more than 8,000, a 16-fold reduction in candidate burden (Fig. 5e). This recovery of unseen clinically supported targets motivated a proteome-wide analysis of candidates beyond those included in the benchmark. ProtScape nominated 102 candidates, approximately half as many as Pinnacle, while achieving around twofold greater enrichment for broader PD associations in Open Targets and approved-drug targets in DrugBank (Fig. 5f). These candidates formed coherent STRING modules with established PD targets and were enriched for synaptic neurotransmission and neuronal excitability, processes central to PD pathophysiology^60,61,62^ (Fig. 5g; Extended Data Fig.5d).

ProtScape also extended established target families towards less-explored mechanisms. It prioritised CACNA1E, CACNA1G, CACNA1H and CACNA1I, which encode R-type CaV2.3 and T-type CaV3.1–CaV3.3 channels, rather than the L-type channels CaV1.2 and CaV1.3 that have dominated calcium-channel therapeutic investigation in PD^63^. Although the L-type channel inhibitor isradipine did not slow clinical progression in a phase 3 trial^64,65^, preclinical evidence implicates CaV2.3 and T-type channels in dopaminergic-neuron vulnerability and pathological firing^66,67,68^. ProtScape therefore nominates alternative calcium-channel subtypes for selective investigation.

Together, these results show that ProtScape improves therapeutic-target prioritisation across diseases while substantially narrowing proteome-wide discovery towards candidates supported by convergent disease, pharmacological, interaction and cellular-context evidence. Its context-resolved attributions further identify the cellular environments most strongly supporting individual predictions, providing a focused basis for experimental target evaluation.

## Discussion

Here we present ProtScape, a multiscale framework that advances context-specific protein modelling by jointly rethinking how cellular interactomes are constructed, learned and evaluated at scale. Rather than treating contextual interactomes as fixed derivatives of reference PPIs and transcriptomic profiles, ProtScape integrates these data to construct biologically grounded interaction scaffolds, learn contextual protein representations, and infer candidate context-specific interactions.

ProtScape substantially improves recovery of held-out Cell-PPI edges while retaining the flexibility to infer interaction landscapes beyond these computational scaffolds. This flexibility is guided towards biologically plausible context-specific interactomes by ProtScape’s architecture, which combines molecular priors, context-dependent topology and scale-adapted modelling. Across reconstruction objectives, stronger Cell-PPI recovery was consistently associated with stronger STRING evidence, and with higher prediction performance for protein-complex membership and therapeutic-target prioritisation across 15 diseases. The consistent advantage of BCE further suggests that closer alignment with the initial Cell-PPI scaffolds strengthens transfer to downstream biological tasks. STRING, however, provides context-independent interaction support and draws partly on the same broader evidence ecosystem as the reference interactomes. Its concordance with ProtScape, therefore, supports the general plausibility of the predicted edges but does not validate their assignment to specific cellular contexts. Matched context-specific interaction measurements will be required to determine which newly inferred edges represent genuine cellular rewiring.

A central promise of context-specific interactome modelling is to move beyond gene expression and capture how proteins are organised within the interaction neighbourhoods that support cellular function. The potential of this approach for disease-mechanism discovery was illustrated in ALS motor neuron differentiation. ProtScape identified predicted stage- and VCP-associated remodelling of the ALS2CL neighbourhood involving RAB-dependent trafficking, despite comparable expression of the implicated proteins. These findings nominate a testable convergence between altered endosomal-network organisation and early synaptic degeneration, illustrating how similar molecular components may be reorganised as cellular state changes. Because differentiation and disease progression are intrinsically dynamic, extending ProtScape beyond discrete contexts towards continuous or temporally resolved graph learning could more directly capture the direction and dynamics of network transitions^69,70^. Experimental studies will also be needed to establish whether these predicted relationships occur physically and contribute causally to the disease phenotype.

Beyond interaction inference, ProtScape’s contextual representations, combined with weak supervision and context-level attribution, provide a framework for resolving protein-level annotations across cellular environments. Protein-complex predictions drew most strongly on contexts containing larger fractions of their constituent proteins and interactions, whereas therapeutic-target predictions highlighted disease-coherent contexts, prioritising immune populations in inflammatory and autoimmune diseases and neuronal populations in Parkinson’s disease. In Parkinson’s disease, ProtScape narrowed the proteome-wide search for clinically supported targets by 16-fold while nominating new candidates embedded in coherent disease-relevant molecular systems. Together, these results show how contextual representations can both focus target prioritisation and identify cellular environments for mechanistic follow-up. Incorporating subcellular localisation, pathway activity and context-resolved proteomics could further refine hypotheses about where protein functions are realised^71,72^. For therapeutic discovery, modelling the hierarchy connecting indications, drugs, interacting proteins and cellular contexts could identify drug–target–context relationships most likely to support clinical activity^73^.

ProtScape provides a foundation for more complete models of context-dependent protein interactions. Integrating proteomic, spatial, structural and post-translational data could complement transcriptome-defined Cell-PPIs by directly informing protein availability, colocalisation, regulatory state and molecular compatibility^74^. Beyond protein-centric networks, extending this framework to multimodal interactomes that include RNA could link post-transcriptional regulation to protein-network reorganisation across cellular contexts^75^. Realising this potential would require coupling ProtScape’s capacity to learn from incomplete supervision with targeted, high-confidence context-resolved measurements that iteratively refine predictions. By connecting scalable computational inference with focused experimental measurement, ProtScape offers a route from prioritising candidate network changes towards experimentally anchored models of molecular reorganisation across cellular states and disease trajectories.

## Methods

### Datasets

#### Protein–protein interaction network

We used the global protein–protein interaction (PPI) network introduced in Pinnacle, defined as the union of BioGRID^76^, the Human Reference Interactome (HuRI)^77^, and the Menche et al. interactome^78^. The resulting reference interactome contained 15,461 proteins and 207,641 interactions.

#### Transcriptomic data

We integrated three transcriptomic resources to capture broad healthy cellular diversity, extend coverage to brain-specific cell types, and include disease-relevant contexts. 1) Tabula Sapiens v1^38^ provided a multi-organ human single-cell reference comprising 15 donors, 59 specimens and 483,152 quality-controlled cells across 24 tissues, excluding the brain. After ontology harmonisation and pseudobulking, 462,841 cells were retained and aggregated into 145 cell-type pseudobulk profiles, corresponding to 143 mapped Cell Ontology classes in the final merged collection. 2) The Human Brain Cell Atlas^39^ complemented this resource with brain-specific cellular populations. We curated 2,115,250 cells from 4 donors, 10 brain regions and 26 Cell Ontology classes. 3) Finally, ALS-relevant bulk RNA-seq data^40,79^ contributed 43 disease-related contexts: 36 motor-neuron profiles spanning three genotypes (control and two disease-associated mutations), two subcellular compartments (cytoplasmic and nuclear) and six time points, together with 7 astrocyte profiles. The combined collection comprised 207 unique cellular contexts.

### Design of context-specific PPI graphs

We constructed a context-specific PPI graph for each cellular context by combining transcriptomic data with the global reference interactome. The goal was to retain genes that were reliably expressed in a context while producing a graph that was sufficiently connected for message passing and distinct across contexts. Compared with Pinnacle, which identifies activated genes by Wilcoxon rank-sum testing against other cells and subsequently retains the largest connected component (LCC) of the corresponding reference-interactome subgraph^31^, ProtScape separates three distinct steps: reliable-expression filtering, context-specific gene ranking and explicit graph-size selection (Extended Data Fig. 1a). While Pinnacle arbitrarily sets a maximum size for the Cell-PPIs, ProtScape instead selects, for each context, the number of top-ranked genes used to construct the graph by balancing interactome coverage and connectivity against redundancy across other cellular contexts.

#### Reliable expression filtering

First, we applied reliably expressed gene (REG) filtering^41^. Each Tabula Sapiens and HBCA context was represented by a single pooled pseudobulk profile, and each ALS context by a single replicate-averaged bulk profile. We therefore fitted a two-component Gaussian mixture model to each context-specific profile using the distribution of log_2_(count + 1) values across genes, and defined a profile-specific expression threshold as the 99th percentile of the low-expression component. We considered a gene reliably expressed in a context if its expression was at or above this threshold.

#### Context-specific ranking

Because REG filtering alone does not distinguish genes that are broadly expressed across many contexts from those preferentially enriched in a specific context, we ranked REGs within each context using a robust one-versus-rest score. Let *x*_*gc*_ denote the average expression of gene *g* in context *c*. For each gene *g* and context *c*, we compared *x*_*gc*_ with the set of average expression values of the same gene across all other contexts,

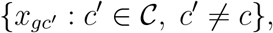

and defined

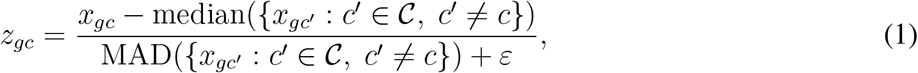

where MAD denotes the median absolute deviation, used with the median for robustness, and *ε* = 10^−6^ is a small constant for numerical stability. For each fixed context *c*, the REGs were ranked in decreasing order of *z*_*gc*_, so that genes with the highest scores were those most specifically enriched in that context relative to the others.

#### Graph-size selection and extraction

We next selected the number of genes retained per context through a data-driven grid search. For each candidate *k*, we took the top-*k* ranked genes for each context, induced the corresponding subgraph of the global PPI, and extracted its LCC. We used the LCC because message passing during pretraining benefits from a well-connected protein graph. For each context *c* and candidate *k*, we computed the LCC coverage |LCC_*c,k*_|*/k*. To quantify cross-context redundancy, we defined the overlap between contexts *c* and *c*^′^ as the Jaccard similarity between their LCC node sets,

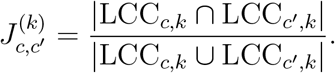

We computed cross-context overlaps with the median across all *c*^′^ ≠ *c* to obtain a robust measure of redundancy with the remaining contexts. For each context, we then selected the value of *k* that maximized

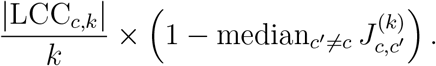

The final context-specific PPI was defined as the LCC of the induced subgraph on the selected genes. We discarded contexts whose final graphs contained fewer than 200 proteins to ensure training stability. This procedure allowed graph size to vary across contexts while retaining both broadly expressed genes needed for connectivity and genes preferentially enriched in specific contexts.

### Design of the metagraph

To connect context-specific PPI graphs across cellular and tissue scales, we constructed a higher-level metagraph whose nodes represent cellular contexts and tissues, and whose edges encode cell–cell interactions (CCIs) and cell–tissue associations.

#### Cell–cell interactions

We inferred cell–cell communication using CellPhoneDB with database v3.0.0^43^, which combines curated ligand–receptor (LR) interactions with a permutation-based statistical framework to identify cell-type-specific LR interactions from gene-expression data. CellPhoneDB evaluates each LR pair by randomly permuting cell labels and comparing the observed interaction mean with the corresponding empirical null distribution.

For the single-cell datasets, we generated CellPhoneDB inputs from the filtered cell-level expression matrices associated with cellular contexts for which Cell-PPIs had been constructed. We sampled up to 100 cells per context before applying 25% cell subsampling. For ALS, we analysed the 43 replicate-averaged, log_2_(count + 1)-transformed context profiles together without subsampling, because each context was represented by a single bulk profile rather than multiple individual cells. We treated each profile as one observation and used the resulting ligand–receptor support to define putative associations between contexts. For all analyses, we performed 100 permutations and retained LR pairs with *P <* 0.001. We then aggregated significant LR pairs at the level of cellular contexts, connecting two contexts by a CCI edge when the number of supported LR pairs reached a dataset-specific threshold.

We selected support thresholds from sensitivity analyses to remove weakly supported CCI edges while preserving coverage of cellular contexts. We used thresholds of 15 significant LR pairs for HBCA and ALS, and 40 for the merged Tabula Sapiens/HBCA graph. At these thresholds, 21 of 22 HBCA contexts, 157 of 168 contexts in the merged Tabula Sapiens/HBCA graph and all 43 ALS contexts retained at least one CCI edge.

We harmonised the dataset-specific CCI graphs using Cell Ontology mappings and combined them by taking the union of edges between matched cellular contexts. We subsequently removed CCI edges involving ALS-derived nuclear-compartment contexts because these contexts represent subcellular fractions rather than whole cells and are therefore not biologically compatible with ligand–receptor-based cell–cell communication. The resulting integrated CCI graph had a density of 0.265, comparable to that of the CCI graph used by Pinnacle (0.293).

#### Cell–tissue assignments

We assigned each cellular context to all tissues where the corresponding cell type was observed in the source metadata, thereby allowing cell types represented across multiple anatomical sites to retain their multi-tissue associations. We harmonised tissue annotations from the different transcriptomic resources to the BRENDA Tissue Ontology^44^. Mapping combined direct ontology matching with manual curation of dataset-specific tissue labels and brain-region annotations that could not be resolved unambiguously through automated matching. For contexts shared across datasets, we retained the union of their mapped tissue annotations.

We used these cell–tissue assignments as multilabel supervision for cellular-context representations, while performing message passing between contexts only over the CCI graph.

### Final Dataset

The resulting collection comprised 207 context-specific PPI networks, including 164 merged single-cell contexts and 43 ALS condition-specific contexts. These networks contained, on average, 5,473 ± 1,621 proteins and 37,916 ± 20,561 edges per network (Extended Data Fig. 1b-d; medians: 5,713 proteins and 35,747 edges). Across all context-specific networks, we identified 15,128 unique proteins, corresponding to 97.8% of the 15,461 proteins in the global reference interactome. At the cellular level, we obtained 5,658 cell–cell interactions and 696 cell–tissue assignments covering 34 directly annotated tissue terms. These terms belonged to an 86-term BTO hierarchy containing 109 tissue–tissue relations, which we did not use for message passing (Extended Data Fig. 1e-f).

### Problem formulation

Based on the context-specific PPI graphs and higher-level cellular relations defined above, we formulate ProtScape as a hierarchical representation-learning problem over a collection of cellular protein interaction graphs coupled through a metagraph.

Let *G* = (*V, E*) denote the global reference interactome, where *V* is the set of proteins and *E* the set of reference protein–protein interactions. Each protein *v* ∈ *V* is associated with a context-independent molecular representation *x*_*v*_ ∈ ℝ^*d*^ obtained from a pretrained protein language model. We denote by *C* = {*c*_1_, …, *c*_*N*_} the set of cellular contexts. For each context *c* ∈ *C*, transcriptomic information defines a context-specific protein set *V*_*c*_ ⊆ *V* and the corresponding induced interaction graph

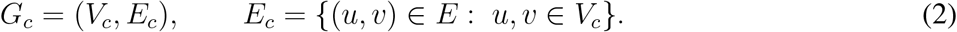

The context-specific graphs are coupled through the cellular graph

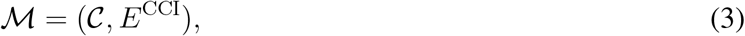

where *E*^CCI^ denotes cell–cell interactions. For each context *c*, the binary vector 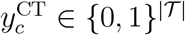 encodes its directly assigned tissues, where *T* is the full 86 BTO tissue vocabulary.

ProtScape aims to transform the same context-independent molecular representation *x*_*v*_ into a context-dependent protein representation 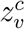 for every occurrence of protein *v* in a Cell-PPI *G*_*c*_. These protein representations are subsequently aggregated into context-level representations that couple the protein and metagraph scales. The learning objectives therefore act at two complementary levels: reconstruction of protein interactions within Cell-PPIs and supervision of higher-order cell–cell interaction reconstruction and cell–tissue multilabel prediction.

### ProtScape architecture

ProtScape used a hierarchical architecture that concentrates representation learning at the protein level while coupling the resulting Cell-PPI representations through higher-level metagraph supervision. All Cell-PPIs were processed by a protein encoder shared across cellular contexts, after which contextualised protein representations were aggregated into context-level embeddings and propagated over the cell–cell interaction graph.

For each context *c*, we considered the undirected graph *G*_*c*_ = (*V*_*c*_, *E*_*c*_) obtained by symmetrising the corresponding Cell-PPI.

#### Initial protein representations

Protein nodes were initialised with context-independent embeddings derived from the sequence-based protein language model ESM-2^33^. For context *c*, the embeddings of proteins in *V*_*c*_ defined the initial feature matrix *X*_*c*_.

To assess the alignment between molecular representations and PPI topology, we quantified node homophily in the global reference interactome using the node-homophily implementation in PyTorch Geometric. Because this analysis required a discrete partition of the continuous ESM2 embedding space, we first evaluated K-means clustering with *K* ∈ {5, 10, …, 40} and then selected the number of clusters based on the silhouette score. This analysis identified *K* = 20 as the optimal partition. ESM2 embeddings were therefore clustered into 20 groups, and node homophily was computed as the mean fraction of neighbours assigned to the same feature cluster. The resulting homophily of 0.151 (or heterophily of 0.849) indicated that interacting proteins predominantly connect proteins occupying distinct regions of the ESM2 representation space.

#### Shared heterophily-aware protein encoder

All context-specific PPIs were encoded using a single GNN backbone with parameters shared across contexts. Consequently, contextualisation resulted from differences in the protein composition and interaction topology of each Cell-PPI rather than from separately parameterised context-specific encoders. This design also avoided the encoder duplication used in Pinnacle and substantially reduces the number of trainable parameters.

The high feature heterophily of the reference interactome motivated the use of Adaptive Channel Mixing (ACM)^34^, instantiated with a random-walk propagation operator. ACM adaptively combined low-pass, high-pass and identity channels, allowing the contribution of neighbourhood smoothing and feature diversification to vary across proteins. Relative to the original ACM formulation, our implementation retained adaptive channel mixing but replaced the GCN-style post-propagation transformation with an MLP-based, GIN-like update^80^, which performed better in our preliminary comparisons than a GCN-style alternative^81^.

Let *A*_*c*_ denote the adjacency matrix of *G*_*c*_, *D*_*c*_ its degree matrix and

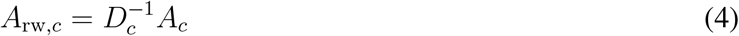

the corresponding random-walk matrix. We define the low-pass, high-pass and identity propagation operators as

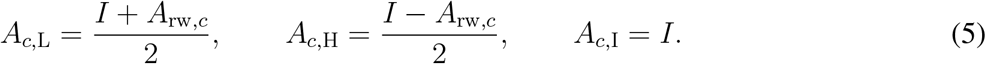

The low- and high-pass operators satisfy *A*_*c*,L_ + *A*_*c*,H_ = *I* and therefore provide complementary views of local neighbourhood information.

Denoting the protein representation matrix at layer *k* by 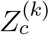, with 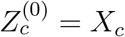, the ACM update is

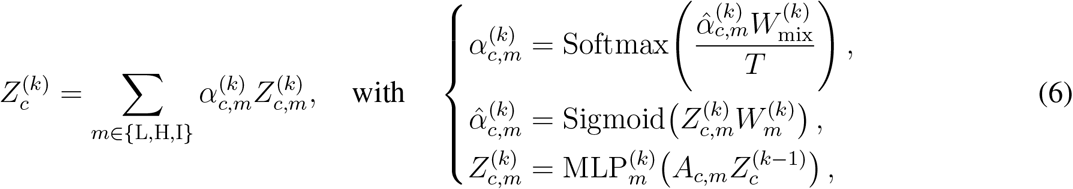

for *m* ∈ {L, H, I}, where *T* is the mixing temperature, 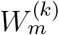 are channel-specific parameters and 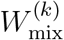 controls mixing across channels. The adaptive weights therefore allow individual proteins to favour smoothing, diversification or retention of their current representation according to their local interaction neighbourhood.

ProtScape used three ACM-RandomWalk layers with a hidden dimension of 512, temperature *T* = 3, leaky ReLU activations, batch normalisation, and dropout. Hyperparameter selection is described below.

#### Multiscale protein representations

To retain information propagated over different neighbourhood depths, we used concatenation-based jumping knowledge^82^. The outputs of the three ACM-RandomWalk layers were concatenated to obtain the final contextualised protein representations

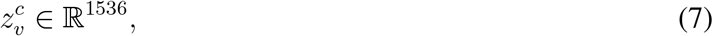

corresponding to 3 × 512 features per protein.

#### Protein-to-context aggregation

Contextualised protein representations were aggregated into a single representation for each cellular context using gated attention pooling^36^. For protein *v* ∈ *V*_*c*_, the attention weight read

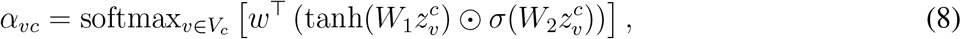

where ⊙ denotes element-wise multiplication, *σ* is the sigmoid function and *W*_1_, *W*_2_ and are *w* learnable parameters. The weights satisfy 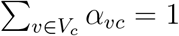. The pooled context representation is

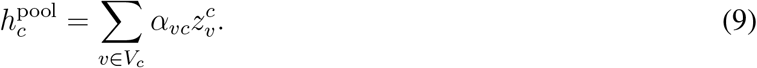

The gated-attention bottleneck has dimension 128.

Because Cell-PPIs were processed in minibatches during training, the representation of a cellular context was estimated from proteins observed across successive updates. We therefore maintained a moving-average memory over 50 updates to stabilise the pooled context representations used by the metagraph-level objectives.

#### Metagraph-level encoding and supervision

The pooled context representations 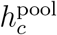 were passed through a single-layer ACM-RandomWalk encoder operating on the training CCI graph, producing CCI-refined context representations 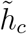. These representations supported two higher-level learning objectives: reconstruction of cell–cell interaction edges and multilabel prediction of the tissues associated with each cellular context.

### Self-supervised pretraining and multiscale learning objectives

ProtScape was trained jointly across the protein and metagraph scales. At the protein level, we compared two self-supervised interaction-reconstruction schemes on the same hierarchical backbone: a standard graph autoencoder (GAE) objective^47^ and a masked graph autoencoding objective based on S2GAE^35^. Both variants used the same shared protein encoder and the same metagraph-level supervision; they differed only in how protein–protein interactions are presented to and reconstructed by the encoder.

#### Standard graph autoencoding

In the GAE setting, protein representations are computed from the observed training graph and used to reconstruct positive protein–protein interactions against structured sampled negatives. Let 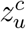 and 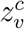 denote the final contextualised representations of proteins *u* and *v* in context *c*. Their interaction score is defined by the dot product

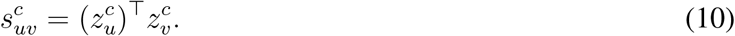

Let 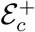 and 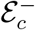 denote the positive and structured negative edge sets for context *c*, and *y*_*uv*_ ∈ {0, 1} the corresponding interaction label. The protein-level reconstruction loss is

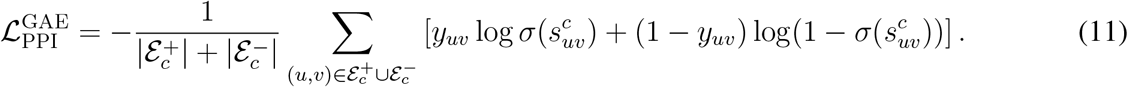

Because the interactions to be reconstructed remain present during message passing, this objective encourages representations that preserve the topology of the observed Cell-PPI.

#### Masked interaction reconstruction

We next considered the S2GAE framework^35^, in which a random subset of positive interactions is removed before message passing and subsequently reconstructed from the remaining graph. For each Cell-PPI, a fraction of positive edges is masked using directed masking; the encoder operates only on the remaining positive interactions, and the masked interactions are predicted against structured sampled negatives. This setting prevents the encoder from directly observing the target interaction and therefore requires it to be reconstructed from molecular representations and the surrounding network topology.

Following S2GAE, let 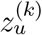 and 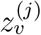 denote the representations of proteins *u* and *v* at encoder layers *k* and *j*. Cross-layer interaction representations are constructed as

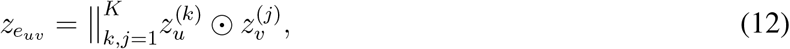

where ⊙ denotes element-wise multiplication and ∥ concatenation. The corresponding interaction logit is predicted using

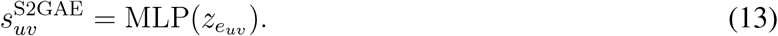

The masked reconstruction loss 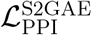 used the same binary cross-entropy formulation as above but was evaluated on masked positive interactions and their associated structured negatives.

#### Metagraph-level supervision

Protein-level learning was complemented by two objectives operating on the CCI-refined context representations 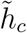. These objectives were identical for the GAE and S2GAE variants, allowing the comparison between the two schemes to isolate the effect of protein-level interaction reconstruction.

Unique undirected CCI pairs were divided into training, validation and test sets using the 0.8*/*0.1*/*0.1 split described above. Only training CCI edges were used for message passing. CCI reconstruction followed an unmasked graph-autoencoding objective in which observed training edges were scored against a 1:1 set of structured negatives, excluding self-loops and all known positive CCI edges. For positive and negative CCI sets 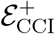 and 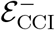, respectively, the metagraph reconstruction loss was

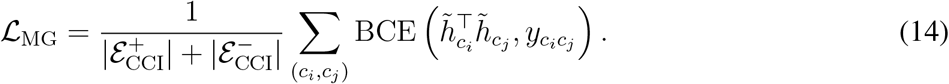

Validation and test CCI edges were excluded from message passing and used only for evaluation.

We used the same CCI-refined representations for multilabel prediction over all 86 tissue terms. Only the 34 directly annotated terms had positive labels and the remaining 52 terms had zero labels across all contexts:

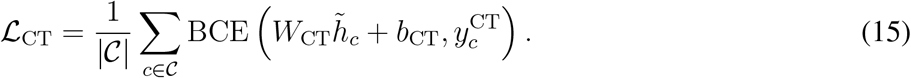

Thus, masked reconstruction is specific to the protein–protein interaction objective, whereas CCI reconstruction uses an unmasked link-prediction objective.

#### Embedding entropy maximisation through uniformity regularization

We additionally monitored whether protein representations retained information across the available embedding dimensions during pretraining. For each trained model, we applied principal component analysis (PCA) to the learned protein representations and quantified the variance explained by each principal component. Let *λ*_*q*_ denote the variance explained by principal component *q* and

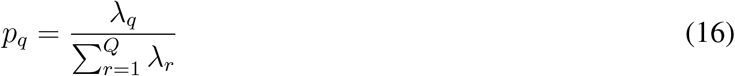

The corresponding normalised explained-variance distribution. We quantified embedding entropy using the normalised Shannon entropy

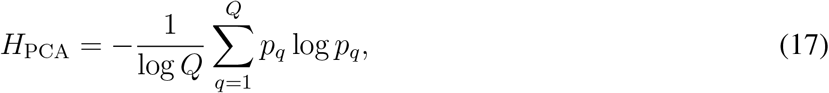

where *Q* is the number of PCA components. *H*_PCA_ approaches one when variance is distributed broadly across components and approaches zero when most variance is concentrated in a small number of directions.

This analysis revealed strong dimensional concentration for the unregularised GAE model, with a normalised PCA entropy of approximately 0.10. We therefore introduced a uniformity regulariser^83^ to encourage the learned protein representations to occupy a broader region of the embedding space. Importantly, the PCA-based entropy was used as a diagnostic and was not itself optimised during training. The implemented regulariser acts on *ℓ*_2_-normalised protein embeddings by minimising the log-average pairwise Gaussian potential,

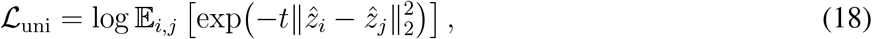

where 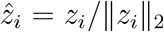 and *t >* 0 is a fixed temperature. Minimising this objective favours a more uniform distribution of representations on the hypersphere, thereby counteracting concentration in a restricted subset of embedding directions.

Masked reconstruction itself substantially alleviated this behaviour: ProtScape-S2GAE without uniformity regularisation reached a normalised PCA entropy of approximately 0.70, which increased further to approximately 0.73 when the uniformity term was included. We therefore use the term *entropy maximisation* to describe the intended effect of this regularisation on representation geometry. In contrast, *uniformity regularisation* refers specifically to the loss implemented above and reported in ablation tables. In ProtScape-S2GAE, adding this term preserved comparable interaction-reconstruction performance at both protein and metagraph scales while improving transfer to downstream tasks, including protein-complex prediction and, most prominently, therapeutic-target prioritisation. Detailed results of ablating this regularisation across our various evaluations are reported in Extended Data Tables 1, 2, and 4.

#### Joint learning objective

The complete ProtScape objective combines protein-level interaction reconstruction, cell–tissue supervision, CCI reconstruction and the optional uniformity term:

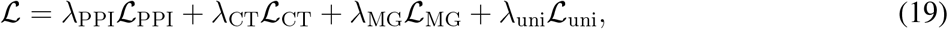

where ℒ_PPI_ denotes either 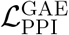 or 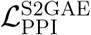. Setting *λ*_uni_ = 0 corresponds to models trained without uniformity regularisation.

Unlike Pinnacle, ProtScape does not explicitly use a centre loss to separate representations of the same protein across cellular contexts. All context-specific occurrences of a protein are initialised from the same molecular representation, while contextualisation emerges from differences in the composition and topology of the corresponding Cell-PPIs. The cell–cell and cell–tissue objectives then organise these representations at the larger metagraph scale. This allows representations of the same protein to diverge when supported by context-specific interaction neighbourhoods without directly imposing separation between contexts.

### Training protocol and model selection

#### Global data splitting and leakage prevention

Because the same protein pair can occur in multiple Cell-PPIs, splitting each context-specific graph independently can assign the same interaction to different data partitions across contexts. In particular, the context-wise splitting strategy used in Pinnacle does not enforce a consistent assignment of recurrent protein pairs across cellular contexts, such that a pair evaluated in one context may already have been observed during training in another. To prevent this form of cross-context leakage, ProtScape uses a single global split defined at the level of unique protein pairs.

We first collected all unique protein pairs occurring across the Cell-PPI atlas and counted the number of cellular contexts in which each pair was present. To preserve comparable proportions of context-specific and recurrent interactions across partitions, recurrence counts were discretised into 10 strata. Unique protein pairs were then assigned to the training, validation, and test sets in the proportions 0.8*/*0.1*/*0.1 using stratified sampling. Each Cell-PPI inherited this global assignment, ensuring that a protein pair assigned to the validation or test set was excluded from training in every cellular context in which it occurred.

The same principle was applied at the metagraph level. Unique undirected CCI pairs were assigned to training, validation and test sets using a 0.8*/*0.1*/*0.1 split, and validation and test interactions were excluded from message passing during training.

#### Mini-batch training

ProtScape models were optimised for 300 epochs using Adam with a learning rate of 10^−2^. Because the complete collection of Cell-PPIs cannot be processed simultaneously, we used GraphSAINT edge sampling^84^ to construct local subgraph minibatches from each context-specific PPI.

GraphSAINT was used exclusively as a graph-sampling mechanism. We did not use its sampling-probability-based node- or edge-reweighting schemes, which produced unstable optimisation when combined with the heterophily-aware ACM encoder in preliminary experiments. Sampled subgraphs were therefore processed directly by the ProtScape encoder without additional GraphSAINT importance reweighting.

The `GraphSAINTEdgeSampler` batch-size parameter was set to 64 and controlled the number of seed edges used to generate each sampled subgraph, rather than the number of complete Cell-PPIs processed jointly. Seed edges determined the sampled protein set, and message passing was performed over the corresponding retained local interactions. At each optimisation step, a subgraph was sampled from each participating Cell-PPI and the resulting context-specific subgraphs were processed jointly. GraphSAINT therefore provided tractable minibatch training while retaining the local PPI topology required by the heterophily-aware message-passing encoder.

#### Negative interaction sampling

Negative protein–protein and cell–cell interactions were generated using structured negative sampling implemented by `structured_negative_sampling` in PyTorch Geometric. For each observed positive interaction, one endpoint was retained, while the other was replaced with a non-neighbour, yielding node-corruption negatives rather than unrestricted, graph-wide random pairs. We adopted this strategy because link-prediction performance is sensitive to negative-sampling design, and uniformly sampled random negatives can produce artificially easy discrimination problems^85^.

Protein-level and CCI reconstruction used one structured negative for each positive interaction, corresponding to a 1:1 positive-to-negative ratio during training. The primary validation and held-out test evaluations used the same 1:1 ratio; evaluations under stronger class imbalance are described separately.

#### Hyperparameter selection

We compared standard GAE and S2GAE protein-reconstruction objectives and varied protein-encoder width in {128, 256, 512}, encoder depth in {3, 4, 5}, dropout in {0, 0.1, 0.2, 0.3, 0.4, 0.5}, protein-to-context aggregation in {mean, attention}, and the cell–tissue loss weight *λ*_CT_ ∈ {0.01, 0.1, 1}. For S2GAE, we additionally evaluated directed and undirected masking, masking ratios in {0.5, 0.7}, decoder widths in {128, 256, 512}, decoder depths in {2, 3} and decoder dropout in {0, 0.2, 0.5}. All candidate configurations were trained using the same global edge split, GraphSAINT sampling protocol and negative-sampling procedure, allowing architectural and objective choices to be compared under identical training conditions. We selected epochs and configurations that maximised the averaged AUPRC across biological scales on the corresponding validation sets.

#### Final ProtScape configuration

The selected ProtScape model used ESM2 protein features and a three-layer ACM-RandomWalk encoder with a hidden dimension of 512, dropout of 0.4, leaky ReLU activations, batch normalisation, and temperature *T* = 3. Concatenation-based jumping knowledge produced 1,536-dimensional contextual protein representations. Protein representations were aggregated using gated attention pooling with bottleneck dimension 128, together with a moving-average context memory over 50 updates.

Protein-level pretraining used S2GAE with directed masking at a ratio of 0.5 and a two-layer cross-layer decoder with width 512, without decoder dropout. The protein-reconstruction, cell–tissue and CCI losses were assigned equal weights, *λ*_PPI_ = *λ*_CT_ = *λ*_MG_ = 1. Uniformity regularisation, introduced above to promote higher embedding entropy, followed Wang and Isola^83^ with *λ*_uni_ = 5 × 10^−5^ and *t* = 2.0. No projection head was used for the uniformity objective.

### Benchmark protocol for context-specific interaction learning

#### Baselines

We used Pinnacle^31^ as the principal baseline for context-specific protein representation learning and retrained all variants on the same Cell-PPI atlas and metagraph used by ProtScape. Pinnacle differs substantially from ProtScape in both parameterisation and information flow. Rather than sharing a single protein encoder across cellular contexts, Pinnacle assigns a distinct GATv2-based GNN to each Cell-PPI. Protein nodes are initialised from high-dimensional random vectors, sampled from a centred and scaled Gaussian distribution, with dimensionality of 1,024 or 2,048 depending on the model configuration.

Pinnacle also does not follow a purely feed-forward protein-to-cell-to-tissue hierarchy. In its first bottom-up pass, each Cell-PPI is processed by two protein-level GNN layers and protein representations are aligned with an attention mechanism before being aggregated into cellular representations through attention-based pooling. The resulting cell embeddings are further updated through residual connections and two GNN layers on the cell–cell interaction graph, followed by non-parametric propagation over tissue–tissue relations. Information is then propagated back from the cellular to the protein scale by unrolling the attention-pooling operation: for protein *v* in context *c*, the context representation contributes proportionally to its previously learned attention weight *α*_*vc*_ through a residual term of the form *α*_*vc*_*h*_*c*_. The updated protein representations are subsequently processed by two additional protein-level GNN layers, followed by a second cell-level aggregation and propagation step. Pinnacle therefore alternates bottom-up and top-down information flow across the metagraph. In contrast, ProtScape uses a shared protein encoder followed by a single forward aggregation towards context- and metagraph-level supervision.

To disentangle the effects of molecular initialisation and graph architecture, we constructed two additional Pinnacle baselines. *Pinnacle–ESM2 (GATv2)* retains the original Pinnacle architecture but replaces its random protein initialisation with pretrained ESM2 representations. *Pinnacle–ESM2 (ACM)* additionally replaces the GATv2 layers operating on the Cell-PPIs and CCI graph with the ACM-RandomWalk layers used by ProtScape, while retaining the remaining Pinnacle information-flow, pooling and supervision scheme. These variants therefore progressively introduce the two main architectural components used by ProtScape—protein language-model features and heterophily-aware message passing—without otherwise replacing the Pinnacle framework.

For direct comparison, all Pinnacle variants were trained on the same 207 Cell-PPIs and metagraph and evaluated using the same global protein-pair splits and structured-negative sampling protocol as ProtScape. All hyperparameters considered in the original Pinnacle paper were validated following the same procedure as for ProtScape models.

#### Held-out edge prediction

Held-out edge prediction followed the global edge splitting and negative-sampling protocol described previously. At validation and test time, held-out positive edges were excluded from message passing and used only for scoring against their corresponding sampled negatives. Validation-time message passing was performed on training edges only, whereas test-time message passing used all non-test edges. For protein-protein interactions, we computed predictions independently in each context and macro-averaged performance across contexts, avoiding metrics skewed by the largest context-specific graphs. We computed CCI performance similarly on held-out test edges. We reported AUPRC and macro-F1 scores for both interaction types.

#### Class-imbalance sensitivity analysis

Cell-PPIs were sparse, with edge densities ranging from 0.16% to 0.50%. We therefore evaluated held-out PPI prediction at 1:*k* positive-to-negative ratios, with *k* ∈ {1, 10, 50, 100, 500}. All models were trained at a 1:1 ratio; larger ratios were used only for evaluation.

### Sensitivity and robustness analysis to incomplete PPI supervision across reconstruction objectives

The Cell-PPIs used for pretraining provide biologically informed but necessarily incomplete supervision. Because they are constructed by selecting context-relevant proteins and inducing the corresponding subgraphs of a global reference interactome, an observed reference interaction may not be active in every cellular context in which both proteins are present. In contrast, biologically plausible interactions absent from the reference interactome are necessarily treated as negatives during reconstruction. We therefore investigated both how strongly ProtScape depends on the assumption that individual Cell-PPI labels should be reproduced exactly and whether variation in predicted interaction strength across cellular contexts reflects reproducible context dependence rather than instability of a particular reconstruction objective.

#### Sensitivity to the PPI reconstruction loss

We trained otherwise identical ProtScape models using three protein-level reconstruction objectives with progressively reduced sensitivity to confident label disagreement: binary cross-entropy (BCE), partially Huberized binary cross-entropy (pHuber) and L1 loss. Architectural components, training splits and higher-level objectives were kept fixed. Dropout was re-evaluated for each reconstruction objective because preliminary experiments showed stronger underfitting with L1. Let *y* ∈ {0, 1} denote the interaction label, *z* the predicted logit and *p* = *σ*(*z*) the corresponding interaction probability.

For BCE,

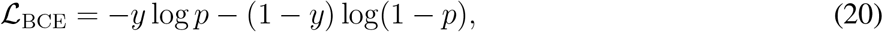

with gradient

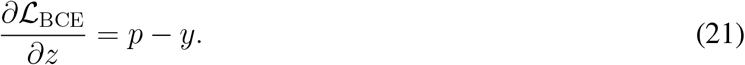

BCE therefore continues to exert a substantial corrective gradient when a confident prediction disagrees with its label. This behaviour is appropriate when supervision is reliable, but gives substantial influence to examples whose supplied labels may be inconsistent with the underlying cellular context.

For L1 reconstruction,

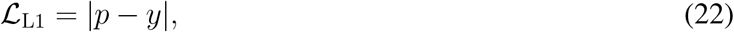

and

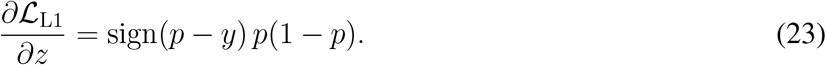

The additional sigmoid derivative reduces the contribution of highly confident predictions, those that disagree with the supplied labels. L1 therefore imposes a weaker requirement that every labelled interaction be reproduced exactly, at the cost of weaker probability calibration and reduced gradients away from the decision boundary.

The pHuber objective^86^ provides an intermediate regime. Denoting by

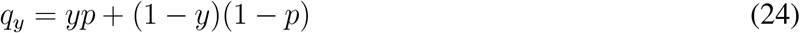

the probability assigned to the observed label, partially Huberized cross-entropy replaces the logarithmic penalty below a transition point with its linear continuation. For transition parameter *τ >* 1,

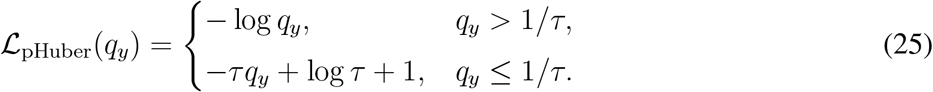

pHuber therefore behaves as BCE for predictions sufficiently compatible with their labels while limiting the influence of extreme disagreements. Together, BCE, pHuber and L1 span increasingly conservative assumptions about the reliability of individual Cell-PPI labels and provide a controlled framework for testing whether ProtScape predictions depend on enforcing the input interaction scaffold.

The selected pHuber configuration used *τ* = 10 and protein-encoder dropout 0.4, matching the dropout of the main BCE model. The selected L1 configuration used no protein-encoder dropout; all remaining architectural and training settings were unchanged.

#### Cross-loss consensus analysis

We next examined whether interaction predictions were reproducible across reconstruction objectives and whether disagreements with the Cell-PPI labels were concentrated in particular regions of prediction space. This analysis was performed post hoc across all 207 Cell-PPIs and independently of the held-out reconstruction benchmark (Extended Data Fig. 2c–i). The BCE-, pHuber-, and L1-trained models were used to score all protein pairs that co-occur within each cellular context. Pairs represented by a Cell-PPI edge were termed labelled positives, whereas co-present pairs without a Cell-PPI edge were termed labelled negatives. The latter, therefore, indicate an absence from the computational interaction scaffold rather than experimentally established non-interactions.

For each interaction–context pair, predictions from the three losses were thresholded at 0.5. Pairs assigned the same negative or positive prediction by all three models were classified as consensus negative (− − −) or consensus positive (+++), respectively. For discordant predictions, the mean prediction determined whether the pair was assigned to the negative or positive side of the consensus, while prediction variability across losses distinguished weak disagreement (− −, ++) from strong disagreement (−, +). The standard-deviation threshold separating weak and strong disagreement was 0.316, corresponding to the mean standard deviation among discordant predictions.

Agreement across losses was concentrated at the extremes of the prediction range, whereas intermediate predictions were more sensitive to the reconstruction objective (Extended Data Fig. 2c,d). This provided a first distinction between robust predictions and uncertainty associated with the choice of reconstruction loss. Importantly, however, loss sensitivity and context dependence are not equivalent: a protein pair can be predicted consistently by all three objectives within a given context, yet change its predicted class across cellular environments. We therefore analysed these two sources of variability separately.

#### Cross-context consistency and contextual variability

For each unique protein pair, we examined its consensus assignments across the cellular contexts in which both proteins were present. Pairs were assigned their most frequent consensus class separately among labelled-positive and labelled-negative occurrences, with ties resolved towards the more negative class. Cross-context stability was assessed for protein pairs represented in at least two contexts. Pairs retaining the same consensus class in every context were considered stable, whereas those assigned different classes across contexts were considered context-variable.

The resulting distributions were strongly structured by consensus strength (Extended Data Fig. 2f,g). Among labelled positives, approximately 157,900 of 196,900 unique interactions (~ 80%) were assigned to the (+ + +) majority class. Conversely, approximately 96 million of 106.6 million labelled-negative pairs (~ 90%) were assigned to the (− − −) class. Extreme consensus classes were also the most stable across cellular contexts, whereas intermediate classes were dominated by pairs whose assignments changed between contexts. Thus, most interactions that were strongly supported or rejected by ProtScape were reproducible across both reconstruction objectives and cellular environments, while contextual variation was concentrated in a smaller subset of the interaction space.

To assess this variation continuously, rather than through consensus classes alone, we also summarised each protein pair by the mean and standard deviation of its ProtScape prediction scores across cellular contexts. The mean captures how broadly an interaction is predicted across cellular environments, whereas the standard deviation captures the extent to which its predicted strength changes between contexts. These quantities were computed independently for the BCE-, pHuber- and L1-trained models and subsequently compared with external STRING evidence.

#### External STRING support for contextual interaction patterns

We used STRING v12.0 to ask whether the observed cross-context prediction patterns were associated with independent graded evidence of interaction. STRING combined scores integrate experimental, computational and literature evidence accumulated across studies and biological settings. ProtScape was trained solely on binary Cell-PPI labels and had no access to the continuous STRING scores.

Protein pairs were mapped to STRING using preferred gene names, retaining the highest combined score when multiple mappings were available. STRING coverage was defined by a combined score greater than zero. Supported pairs were further grouped into low (0 *< s <* 0.4), intermediate (0.4 ≤ *s <* 0.8) and high (*s* ≥ 0.8) evidence categories. Because STRING does not resolve interactions by cellular context, it was not used as a direct ground truth for context-specific predictions. Instead, we tested whether global STRING evidence varied systematically with the overall strength and contextual variability of ProtScape predictions. We used ordinary least squares to predict STRING combined scores from the across-context mean and standard deviation of ProtScape predictions, separately for each loss and jointly across losses. We fitted the regressions on 80% of STRING-supported protein pairs co-present in at least two contexts and evaluated Pearson and Spearman correlations on the remaining 20%.

Approximately 60% of labelled-positive reference interactions were represented in STRING (Extended Data Fig. 2h). STRING coverage itself remained relatively similar across the labelled-positive consensus classes. Still, the distribution of continuous STRING scores shifted systematically with ProtScape predictions: interactions assigned to increasingly positive consensus classes showed progressively stronger STRING evidence, whereas those assigned towards the negative side showed weaker support (Extended Data Fig. 2i). The same pattern was recovered in the continuous analysis reported in the main text. Across reconstruction objectives, STRING scores increased with the mean ProtScape prediction across contexts and decreased as cross-context prediction variability increased. Protein pairs predicted strongly and consistently across many cellular environments therefore tended to have greater context-independent STRING support, whereas interactions with more context-restricted prediction profiles accumulated weaker global evidence.

This relationship is consistent with the way STRING evidence is assembled. Interactions recurrently observed across experimental systems and biological settings are more likely to accumulate multiple sources of support and consequently high combined scores. In contrast, interactions restricted to specific cellular environments may be less well documented. The observed inverse relationship between STRING evidence and cross-context variability therefore supports the interpretation that variation in ProtScape predictions across cellular contexts is structured rather than arbitrary, while not implying that STRING directly validates the cellular specificity of individual interactions.

The analysis of labelled negatives provided an additional test of incomplete supervision. Although most labelled-negative pairs were assigned to the (− − −) class, approximately 1.2 million of 106.6 million unique labelled-negative pairs (~1.1%) received a majority consensus-positive (+ + +) assignment across contexts despite being absent from the Cell-PPI scaffold. Approximately 204,000 of these interactions were represented in STRING, corresponding to a coverage of ~ 17%, compared with only ~ 4.2% among consensus-negative (− − −) labelled pairs. Consensus-positive labelled negatives were therefore approximately fourfold enriched for STRING-supported interactions. Intermediate consensus classes showed a corresponding gradient of STRING coverage and evidence strength from the positive to the negative side.

These findings indicate that disagreement with the Cell-PPI labels cannot be interpreted uniformly as reconstruction error. A small but substantial subset of pairs absent from the computational scaffold is reproducibly predicted across reconstruction objectives and independently supported by STRING, consistent with false-negative supervision arising from incompleteness of the reference interactome. At the same time, the strong concentration of labelled positives in the (+ + +) class and labelled negatives in the (− − −) class shows that ProtScape remains predominantly anchored to the initial Cell-PPI structure.

Taken together, the cross-loss, cross-context and STRING analyses provide complementary evidence for the structure of the inferred interaction landscapes. Predictions that are stable across reconstruction objectives distinguish robust model behaviour from loss-specific uncertainty. In contrast, variation of the same protein pair across cellular environments captures context-dependent changes in inferred interaction strength. The systematic increase in STRING evidence with prediction strength, together with its decrease as contextual variability increases, supports the interpretation that broadly recurrent interactions receive stronger global evidence. In contrast, more context-restricted predictions are less well represented in context-independent interaction resources. The enrichment of STRING-supported interactions among consensus-positive labelled negatives further shows that departures from the initial Cell-PPI scaffold can recover plausible interactions missing from the supervision.

### Weakly supervised downstream learning and post-hoc explainability

Most downstream annotations considered in this study, including protein-complex membership and therapeutic-target labels, are defined at the protein level rather than for individual cellular contexts. We therefore formulated downstream prediction as a weakly supervised multiple-instance learning (MIL) problem, in which each protein is represented by the collection of its context-specific representations and receives a single protein-level label.

This formulation differs from the downstream strategy used in Pinnacle^31^, where the same protein-level label is assigned independently to every contextualised representation of that protein. Such label replication implicitly assumes that a protein annotation is equally valid across all cellular contexts in which the protein is represented. For context-dependent biological functions, this assumption introduces systematic label noise: only a subset of cellular environments may support the annotated function or disease association. Importantly, the same issue also affects evaluation, because each context-specific representation is treated as an independently labelled example even though the available ground truth is defined only at the protein level. Consequently, context-aware models may be penalised for assigning low scores to biologically irrelevant contexts, and performance can depend on the number and distribution of contexts associated with each protein.

To avoid this mismatch, ProtScape assigns supervision only at the level at which annotations are available. Each protein is represented as a bag containing all of its context-specific instances, and the model produces a single prediction after aggregating information across the bag. This allows the downstream model to identify which cellular contexts contribute to a protein-level annotation without assuming that the annotation holds uniformly across contexts, while providing a common protein-level evaluation framework for contextual and context-free representations.

For each downstream task, labelled proteins were restricted to those represented in the global interactome. A held-out test set was first defined, after which five-fold cross-validation was performed on the remaining proteins for model selection. Splits were stratified to preserve the distribution of labels and contextual representation patterns across folds.

#### Context-free and mean-pooled baselines

We first evaluated context-free protein representations using logistic-regression probes trained on ProstT5 or ESM2 embeddings. Contextual representations were additionally evaluated using mean pooling, in which all context-specific instances associated with a protein contribute equally to the resulting bag representation. Linear classifiers were trained either on these pooled contextual features alone or after concatenation with the corresponding context-free ESM2 embedding. These models provide a non-attentive MIL baseline in which no cellular context is preferentially weighted.

#### Attention-based multiple-instance learning

For protein *i*, let

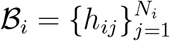

denote the bag of its *N*_*i*_ context-specific instances. Each instance *h*_*ij*_ is formed by concatenating the contextualised protein representation with the representation of the corresponding cellular context. Let *e*_*i*_ denote the context-free sequence embedding of protein *i*.

We used gated attention-based multiple-instance learning (ABMIL)^36^ to learn the contribution of each cellular context to the protein-level prediction. Rather than concatenating the sequence embedding to every contextual instance, we adopted a late-fusion strategy: attention is computed exclusively from the contextual instances, and the context-free protein representation is introduced only after bag aggregation. This preserves a context-specific attention mechanism while allowing the sequence embedding to act as a global molecular prior. Replicating the same sequence representation across all instances would otherwise introduce an identical component into every attention input, potentially reducing the contrast between cellular contexts.

For contextual instance *j*, the attention score is

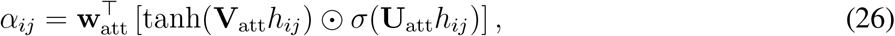

and the corresponding normalised attention weight is

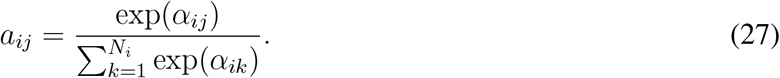

The contextual bag representation is then

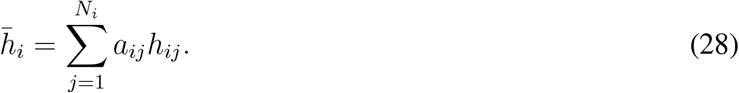

When sequence late fusion is used, the final prediction is obtained as

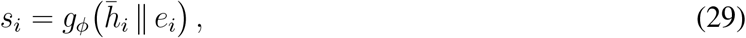

where *g*_*ϕ*_ denotes the task-specific classifier and ∥ concatenation. Without sequence late fusion, *g*_*ϕ*_ operates directly on 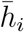. For multi-head variants, eight independent attention heads were used, and their pooled contextual representations were concatenated before classification.

#### Progressive regularisation of ABMIL

The downstream datasets considered here are generally small relative to the dimensionality and number of contextual instances available for each protein, making attention-based readouts susceptible to overfitting. We therefore used progressive dropout layers (PDL)^87^ as the default regularisation strategy for ABMIL.

PDL combines an importance-dependent within-bag dropout mechanism with a progressive training schedule. Contextual instances are first passed through a same-dimensional two-layer nonlinear transformation containing PDL modules. Within each bag, instances are ranked by learned importance scores, with lower-ranked instances assigned higher dropout probabilities. The maximum dropout probability is progressively increased during training, while retaining at least one instance per bag. This encourages the classifier to distribute evidence across multiple informative contexts rather than relying prematurely on a small number of high-weight instances.

Unless explicitly stated otherwise, *ABMIL* in the main text refers to this PDL-regularised gated-attention model. We report the corresponding unregularised gated-attention architecture separately in the supplementary ablations as ABMIL without PDL, whereas we denote the regularised implementation as ABMIL–PDL in the Extended Data Tables for clarity. We selected ABMIL–PDL as the default downstream readout because it consistently generalised better on the relatively small labelled datasets considered in this study.

#### Training and model selection

We froze all pretrained ProtScape and Pinnacle protein and context representations during downstream evaluation; we optimised only the task-specific readouts. Models were trained with AdamW using a learning rate of 10^−4^, weight decay 10^−4^, batch size 512 and a maximum of 300 epochs. We used early stopping on validation AUPRC with patience 50.

We performed feature normalisation independently within each training fold to prevent information leakage. Z-score statistics for contextual representations were estimated using all instances belonging to training proteins only, whereas sequence-embedding statistics were estimated from the corresponding training proteins. We then applied the resulting parameters unchanged to validation and test proteins.

We constructed training batches using a multilabel-stratified sampler based on precomputed label clusters to preserve class composition across minibatches. All downstream tasks used class-weighted binary cross-entropy with logits. We selected hyperparameters and checkpoints based exclusively on validation AUPRC within each fold, then evaluated the resulting models on the fixed held-out test set. Reported test performance corresponds to the mean across the five fold-specific models.

#### Post-hoc interpretation with layer-wise relevance propagation

We interpreted the trained ABMIL models using xMIL-LRP^**?**^, a MIL-specific layer-wise relevance propagation that provides signed, class-specific attributions. We computed explanations after training, with all model parameters fixed.

For protein *i* and output class *t*, the pre-sigmoid logit *s*_*it*_ was used as the explanation target. Relevance was propagated separately to the contextual bag 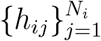 and, when present, to the late-fused sequence representation *e*_*i*_. This produced feature-level relevance scores 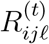 for feature *ℓ* of contextual instance *j*, and 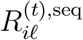 for the sequence branch.

Context-level signed relevance was obtained by summing over embedding dimensions,

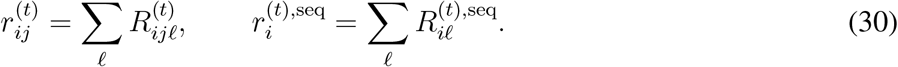

Positive relevance denotes evidence supporting class *t*, whereas negative relevance denotes evidence opposing it.

To quantify the relative contribution of contextual and sequence information, we computed the fraction of absolute feature-level relevance assigned to the contextual branch:

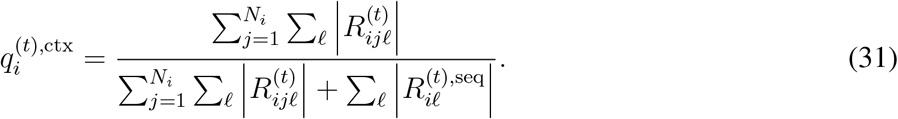

Taking absolute values before aggregation prevents positive and negative relevance from cancelling. The resulting score ranges from 0 to 1, with larger values indicating greater reliance on contextual representations relative to the sequence prior.

For analyses requiring attribution to individual cellular contexts, including the protein-complex and therapeutic-target analyses, we focused on positive contextual support. For context *j*,

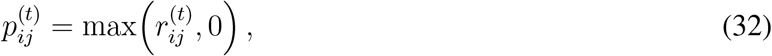

and normalised this contribution relative to all positive evidence from the contextual and sequence branches:

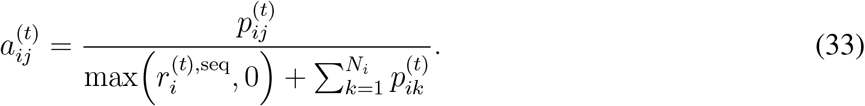

If the denominator was zero, all normalised contextual contributions were set to zero. Because positive sequence relevance is retained in the denominator, the contextual contributions need not sum to one. Only cellular contexts represented in a protein’s bag were included; absent contexts were not assigned zero-valued pseudo-instances. We calculated relevance scores independently for each fold-specific model before aggregating across folds.

### CORUM protein-complex prediction and contextual interpretation

The CORUM benchmark was derived from human protein-complex membership in CORUM v4.1^51^. We mapped complex subunits to HGNC gene symbols and restricted them to proteins represented in the global interactome. We retained complexes with 10–300 proteins and removed highly redundant entries when either their overlap coefficient was at least 0.90 or their Jaccard similarity was at least 0.80. After processing, we retained 155 complexes, 1,332 proteins and 2,414 protein–complex memberships. We restricted evaluation to proteins with both contextual and ESM2 embeddings, retaining 1,328 proteins and 2,409 memberships across all 155 complexes. Retained complexes contained 10–104 proteins, with a median size of 12. Because individual proteins can belong to multiple complexes, we formulated prediction as a multilabel classification problem.

CORUM annotations are defined at the protein level and do not specify the cellular contexts in which individual complexes assemble or function. We therefore used the weakly supervised framework described above rather than assigning each complex-membership label independently to every contextualised protein representation. Unless otherwise stated, contextual models used the default ABMIL-PDL readout with ESM2 late fusion, referred to as ABMIL in the main text. We evaluated sequence-only and mean-pooled contextual representations as complementary baselines, and kept all pretrained representations frozen.

The benchmark followed the downstream evaluation protocol described above. We partitioned proteins into six multilabel-stratified folds, retained one fold of 222 proteins as a fixed test set, and used the remaining five iteratively for training and validation. We selected models based only on validation AUPRC, then evaluated each fold-specific model on the same held-out test set. The reported AUPRC and F1 values are macro-averaged across complexes, then averaged across the five selected models. We used the same protocol for the reconstruction-loss sensitivity analysis.

#### Protein-complex structural analyses

To determine which protein-complex properties were associated with prediction performance, we considered its size, its representation across Cell-PPIs, and its connectivity in the global reference interactome. Let *Q* denote the set of proteins belonging to a CORUM complex, *G* = (*V, E*) the undirected global interactome and *G*_*c*_ = (*V*_*c*_, *E*_*c*_) the Cell-PPI associated with context *c*. The set of global-interactome edges induced by *Q* was defined as

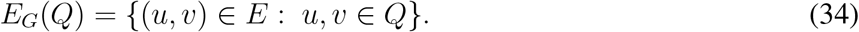

For each context, complex node coverage was defined as

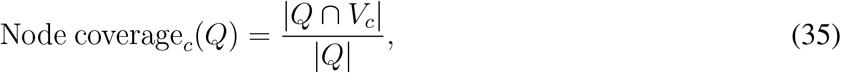

and complex edge coverage as

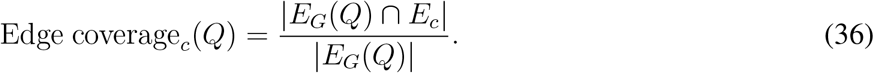

These quantities measure, respectively, the fraction of complex members and the fraction of global-interactome complex edges retained in a given Cell-PPI. We averaged both across all 207 cellular contexts, including contexts in which no complex member or interaction was represented.

Connectivity of the complex in the global interactome was quantified independently of cellular context as

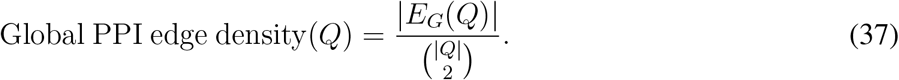

For each complex, we first calculated average precision independently for each of the five fold-specific models, then averaged across models. Associations between complex-level performance and each structural descriptor were assessed across the 155 complexes using two-sided Spearman rank correlations.

For visualisation, we ranked complexes independently for each descriptor and divided them into 10 equal-count percentile bins. Binning was used only for display; all correlation coefficients and *P* values were calculated from the original unbinned values.

#### Context-level interpretation of CORUM predictions

We used xMIL-LRP, as described above, to test whether ABMIL preferentially relied on cellular contexts in which a predicted complex was more completely represented. Explanations were computed using the validation-selected ABMIL-PDL models with ESM2 late fusion and were restricted to held-out proteins positively annotated to the corresponding complex.

For protein *i*, complex *Q* and cellular context *c*, we first calculated the normalized positive contextual relevance 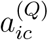 defined above. This quantity measures the fraction of the complete positive attribution assigned to context *c*, while retaining any positive contribution from the late-fused sequence branch in the normalisation denominator.

To obtain a single attribution score for each complex–context pair, 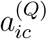 was first calculated independently for each of the five fold-specific models and averaged across models. The resulting values were then averaged across held-out proteins positively annotated to complex *Q* and represented in context *c*:

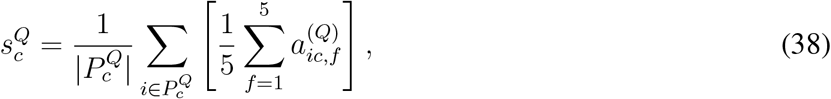

where 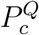 denotes the set of positively annotated held-out proteins from complex *Q* whose bag contains context *c*, and *f* indexes the five fold-specific models. We excluded contexts absent from a protein’s bag rather than assigning them a relevance of 0.

For each complex, we then assessed whether contexts receiving greater positive attribution also contained a larger fraction of the corresponding complex. This was quantified separately for node and edge coverage using

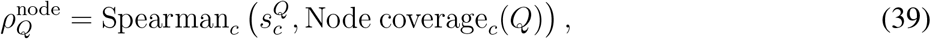

and

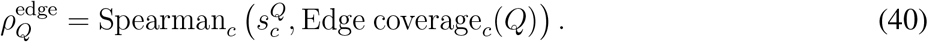

We calculated correlations only when at least three eligible contexts were available, and the corresponding coverage measure varied across contexts. When contextual relevance was constant, we set the correlation to zero. The median correlation across complexes was reported separately for node and edge coverage.

For visualisation, we divided complex–context pairs into 10 equal-count percentile bins based on node or edge coverage. Within each complex, contextual relevance 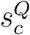 was first averaged across contexts in the same bin, and these complex-level values were subsequently averaged across complexes. As above, we used binning only for visualisation, and performed all correlation analyses on unbinned complex–context observations.

### ALS-associated network rewiring

#### Late-differentiation interaction networks

We analysed motor-neuron Cell-PPIs at days 22 and 35 of differentiation in control (CTRL) and VCP-mutant (VCP) conditions, considering nuclear and cytoplasmic compartments separately. Protein pairs represented in at least one compartment for either genotype at either time point were retained and scored using the trained S2GAE model.

Protein pairs absent from the global reference interactome were considered high-confidence newly inferred interactions when their S2GAE score was ≥ 0.95 in at least one compartment. At each time point, newly inferred interactions detected in only one genotype were classified as condition-specific, yielding four sets: CTRL-D22, VCP-D22, CTRL-D35 and VCP-D35. The resulting ALS-associated network comprised all newly inferred, condition-specific interactions, together with all reference interactions connecting proteins incident to at least one such edge.

#### Module detection and refinement

We identified network communities using the Leiden algorithm with the RBConfiguration quality function. We evaluated the resolution parameter over the range 0.25–2.00 and selected it by jointly considering modularity and partition stability across 10 random seeds. We generated the final partition at the selected resolution using seed 42.

To resolve large heterogeneous communities, we subsequently refined the partition by iteratively re-clustering the least internally connected module. Refinement was stopped at the elbow of the mean intra-module connectivity curve, yielding a final partition of 80 modules (Extended Data Fig. 4a).

To assess whether higher structural resolution also improved functional separation between modules, we represented each module in Gene Ontology (GO) pathway space using a size-normalised pathway-membership score. For module *M* and GO term *g*, the score was defined as

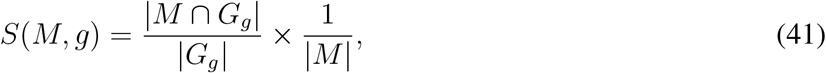

where *G*_*g*_ denotes the set of genes annotated to GO term *g*. We quantified functional dissimilarity between modules by Euclidean distance between their pathway embeddings. We evaluated changes in functional separation alongside modularity and intra-module connectivity during successive refinement steps (Extended Data Fig. 4a).

#### Enrichment of genotype-specific rewiring across modules

For each module and time point, we tested whether newly inferred CTRL- and VCP-specific interactions were distributed differently from the remainder of the network. Counts of genotype-specific interactions inside and outside each module were compared using two-sided Fisher’s exact tests, with *P* values adjusted separately at days 22 and 35 using the Benjamini–Hochberg procedure.

Fold enrichment was defined as the fraction of all VCP-specific interactions assigned to a module divided by the corresponding fraction of CTRL-specific interactions. Modules were considered genotype-enriched when they showed both a minimum 20% shift in relative edge representation and an FDR below 0.01; VCP-enriched modules therefore had fold enrichment *>* 1.2, whereas CTRL-enriched modules had fold enrichment *<* 0.83 (Extended Data Fig. 4b).

#### Functional enrichment analysis

We tested genes in modules of interest for enrichment of Reactome pathways and Gene Ontology terms using the STRING functional enrichment API. Functional categories were evaluated using the FDR-corrected enrichment statistics returned by STRING.

#### Gene-expression analysis

Filtered normalised gene-expression values across differentiation (days 0–35) were transformed as log_2_(*x* + 1) and averaged across replicates. Expression dynamics were then expressed as log_2_ fold changes relative to the corresponding genotype-specific day-0 baseline. We used cytoplasmic RNA measurements as a proxy for gene expression.

Because the same genes were compared between CTRL and VCP conditions, we assessed genotype-associated expression differences within selected modules using paired Wilcoxon signed-rank tests. This analysis determined whether condition-specific interaction rewiring could be explained by corresponding changes in transcript abundance.

### Therapeutic-target prediction and Parkinson’s disease prioritisation

#### Therapeutic-target benchmark

We evaluated 15 disease-specific therapeutic-target prediction tasks adapted from the Pinnacle benchmark^31^. The indications comprised inflammatory bowel disease, rheumatoid arthritis, breast carcinoma, systolic heart failure, lung adenocarcinoma, psoriasis, pulmonary arterial hypertension, type 2 diabetes mellitus, systemic lupus erythematosus, atopic eczema, asthma, chronic obstructive pulmonary disease, non-alcoholic steatohepatitis, chronic kidney disease and Parkinson’s disease.

We defined positive examples using Open Targets 24.03 ChEMBL clinical evidence and Open Targets 26.03 disease hierarchy, association and target mappings^88^. For each disease, we retained proteins targeted by drugs supported by completed phase II or stronger clinical evidence, including evidence for descendant disease terms. We drew negative examples from the October 2022 DrugBank set of approved-human drug targets^89^, excluding positive targets and proteins with non-literature Open Targets associations to the queried disease. We restricted both classes to the global interactome and provide the frozen processed labels with the data release.

We trained and evaluated all models using the weakly supervised protocol described above, restricting the benchmark to proteins with both contextual and ESM2 embeddings. We report disease-specific cohort sizes in Extended Data Table 3.

#### Context-level interpretation

We interpreted therapeutic-target predictions using the xMIL-LRP framework defined above. Explanations targeted the pre-sigmoid disease logit and were restricted to held-out positive proteins. For protein *i*, disease *t* and context *j*, we used the normalised positive contextual relevance 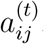.

For cell-type analyses, we summed contextual relevance across contexts that map to the same cell type. For broad cell-class analyses, we instead averaged relevance over the contexts belonging to class *g*,

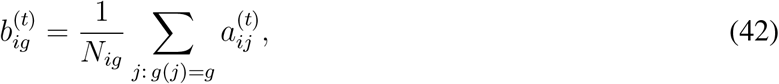

where *N*_*ig*_ is the number of contexts from class *g* represented in the bag of protein *i*. This prevents classes represented by more cellular contexts from receiving larger attribution solely because of their size.

We first averaged scores across the five fold-specific models for each protein, then averaged across held-out positive proteins within each disease. Explanations with no positive contextual relevance contributed zero. For heatmap visualisation, we z-score standardised cell-class scores across classes within each disease. Relative reliance on contextual versus sequence information was quantified separately using 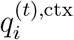, as defined above.

#### Proteome-wide Parkinson’s disease prioritization

We applied the five validation-selected Parkinson’s disease ABMIL models with ESM2 late fusion to all 14,782 proteins for which both a contextual bag and an ESM2 embedding were available. We averaged predictions across the five models to obtain one ensemble score per protein.

For held-out target recovery, we removed all training and validation benchmark proteins and held-out test negatives, leaving 18 held-out positive targets and 13,303 proteins without benchmark labels. We ranked proteins by decreasing ensemble score, and defined screening burden as the number of label-excluded proteins ranked ahead of each recovered held-out target. This provided a threshold-independent measure of the proteome depth required to recover clinically supported targets.

For candidate nomination, we removed all benchmark-labelled proteins and retained label-excluded proteins with ensemble score ≥ 0.5, yielding 102 ProtScape and 205 Pinnacle candidates. We used the threshold as a fixed nomination rule rather than a calibrated operating point. For the kainate-receptor (GRIK1–GRIK5), neuroligin (NLGN1, NLGN2, NLGN3 and NLGN4X) and DLGAP (DLGAP1–DLGAP4) families, completion depth was defined as the lowest-ranking position required to recover all family members.

#### External annotation of Parkinson’s disease candidates

Candidate annotations were obtained from Open Targets queries performed on 28–29 July 2026^88^ and the October 2022 DrugBank collection of approved human drug targets^89^. A current Parkinson’s disease association required a non-literature Open Targets association within the disease hierarchy rooted at MONDO:0005180. Approved-drug support indicated targeting by an approved human drug for any indication, whereas other-disease support denoted an Open Targets disease association in the absence of a retained Parkinson’s disease association. Analyses were performed either on candidates nominated at the 0.5 threshold or on the top *k* label-excluded proteins.

#### STRING network and functional analysis

Benchmark-positive Parkinson’s disease targets and ProtScape candidates nominated at ensemble score ≥ 0.5 were mapped to STRING v12.0^50^. We retained interactions with non-zero experimental evidence and weighted them by their experimental score.

Communities were identified using weighted Leiden clustering with the RBConfiguration quality function^90^. We evaluated resolutions from 0.25 to 2.0 in increments of 0.25 across 10 random seeds. We discarded resolutions that produced a single community in any run. Mean weighted modularity and mean pairwise adjusted Rand index were independently min–max normalised and combined with equal weight to select the resolution; the most stable partition at that resolution was retained.

We assessed the significance of the observed modular structure against 1,000 connected degree-preserving double-edge-swap null networks with permuted edge weights. For the observed network and each null network, we retained the maximum modularity over five Leiden restarts and calculated empirical *P* values using a plus-one correction.

We annotated communities by one-sided hypergeometric enrichment relative to the displayed network. Terms represented by fewer than three proteins or more than 80% of network proteins were excluded, and *P* values were adjusted within annotation source using the Benjamini–Hochberg procedure. We retained the five most significant Reactome terms for each community.

For complementary functional analyses, we considered STRING annotations from GO biological process, GO molecular function, GO cellular component, Reactome, KEGG, Pfam, SMART, and InterPro. Benchmark-positive targets were tested against all mapped benchmark-labelled proteins, whereas candidate sets were tested against all mapped label-excluded proteins. Terms containing 10–500 background proteins were evaluated using one-sided hypergeometric tests with Benjamini–Hochberg correction within each annotation source. Fold enrichment was defined as

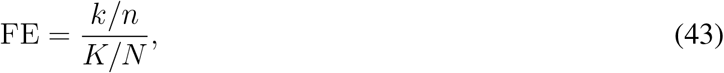

where *k* and *n* denote the number of query hits and query size, and *K* and *N* the corresponding background quantities. For Fig. 5, we selected the three lowest-FDR benchmark-target terms independently from GO biological process, GO molecular function, and Pfam, and displayed the corresponding terms for the ProtScape candidates.

## Data availability

The processed datasets, contextual protein and cell embeddings, and pretrained checkpoints used in this study are available on Zenodo at https://zenodo.org/records/22645081.

## Code availability

All methods developed and used in this study are available on Github at https://github.com/AI-for-RNA-Biology/ProtScape.

### Acknowledgements

This work was financially supported by CRTOH-GTO award no. 2024_SA_24_005 (to L.F.); the Swiss National Science Foundation | National Centre of Competence in Research (NCCR) on RNA & Disease Phase 3 Ref Number: 51NF40-205601 (to C.V.C.; R.L.). This work was supported by a donation to the GTO programme; the Ligue genevoise contre le cancer (LGC) and the Fondation privée of the Geneva University Hospitals. R.P. gratefully acknowledges generous support from NUS (Start up grant), a Lister Research Prize Fellowship, Steve Redgwell, Liane Iles, MND Foundation, the Motor Neuron Disease Association (Patani/Dec22/957-793), My Name’5 Doddie Foundation (MN5DF/2022/003), and Target ALS (BB-2024-C4-L4).

## Author contributions

Conceptualization, C.V.C., R.L.; Methodology, A.T., C.V.C.; Software, A.T., C.V.C., L.F.; Formal Analysis, A.T., L.F.; Investigation, A.T., L.F., V.J., C.V.C.; Writing – Original Draft, C.V.C., R.L., A.T.; Writing – Review & Editing, R.L., C.V.C., A.T., L.F., R.P., P.F.; Resources, R.L., P.F.; Visualization, A.T., C.V.C., R.L., L.F.; Funding Acquisition, R.L., P.F.; Supervision, R.L., C.V.C.

## Extended Data Figures and Tables

**Extended Data Fig. 1.**
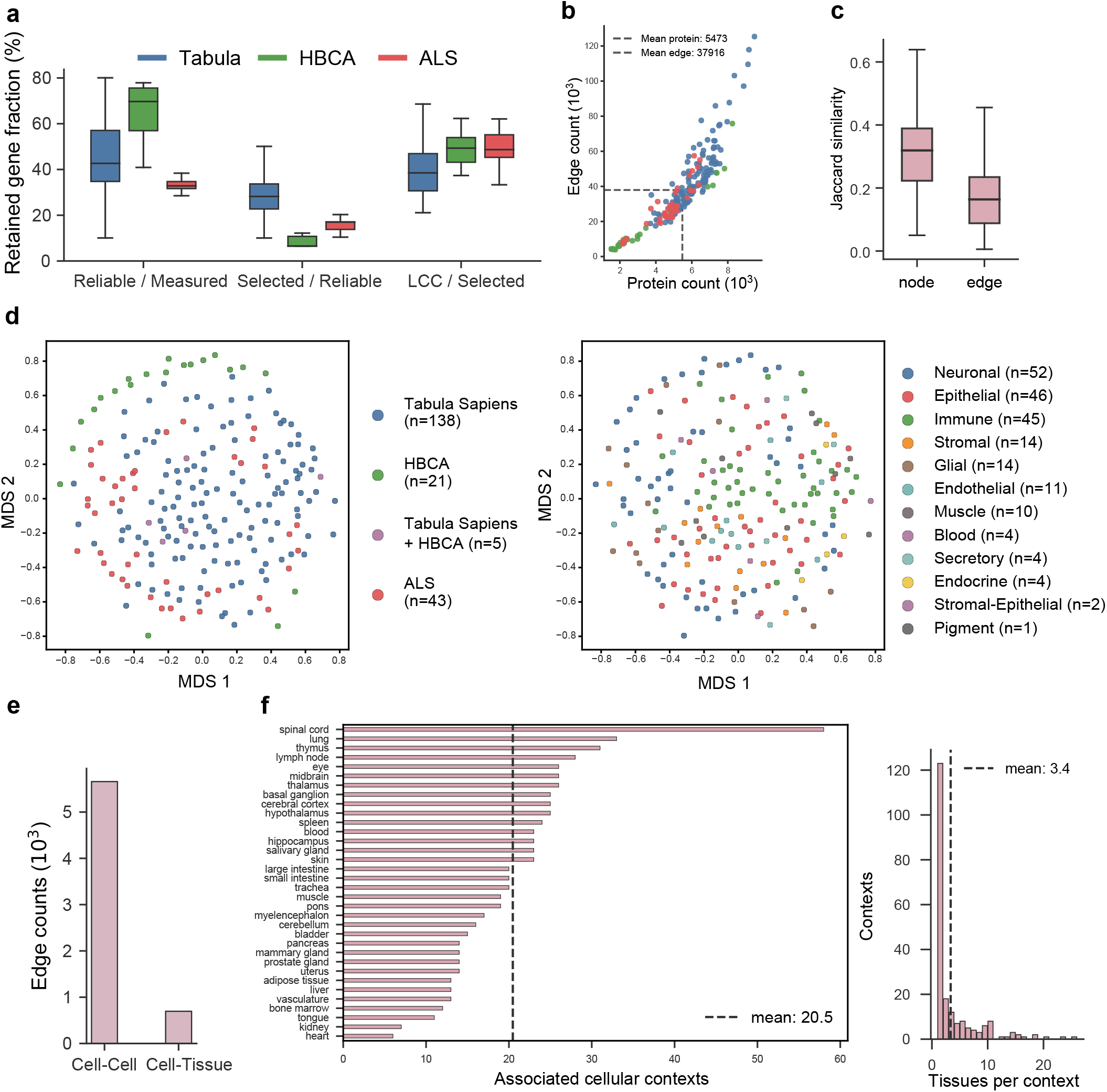
Construction and characteristics of the context-specific interactome atlas and metagraph. **a**, Gene retention across successive stages of Cell-PPI construction for Tabula Sapiens, HBCA and ALS contexts: reliably expressed genes relative to measured genes, context-enriched genes relative to reliably expressed genes, and genes retained in the largest connected component (LCC) relative to selected genes. **b**, Numbers of proteins and interactions in the resulting Cell-PPIs. Each point represents one cellular context and colours indicate transcriptomic source; dashed lines denote mean values across contexts. **c**, Pairwise Jaccard similarity between Cell-PPIs based on retained proteins (nodes) or interactions (edges). **d**, Multidimensional scaling (MDS) of Cell-PPIs based on node-level Jaccard similarity, coloured by transcriptomic source (left) or broad cell class (right). **e**, Numbers of cell–cell and cell–tissue edges in the multiscale metagraph. **f**, Tissue coverage of the Cell-PPI atlas. Left, number of cellular contexts associated with each tissue; right, number of tissues associated with each cellular context. Dashed lines indicate mean values.

**Extended Data Fig. 2.**
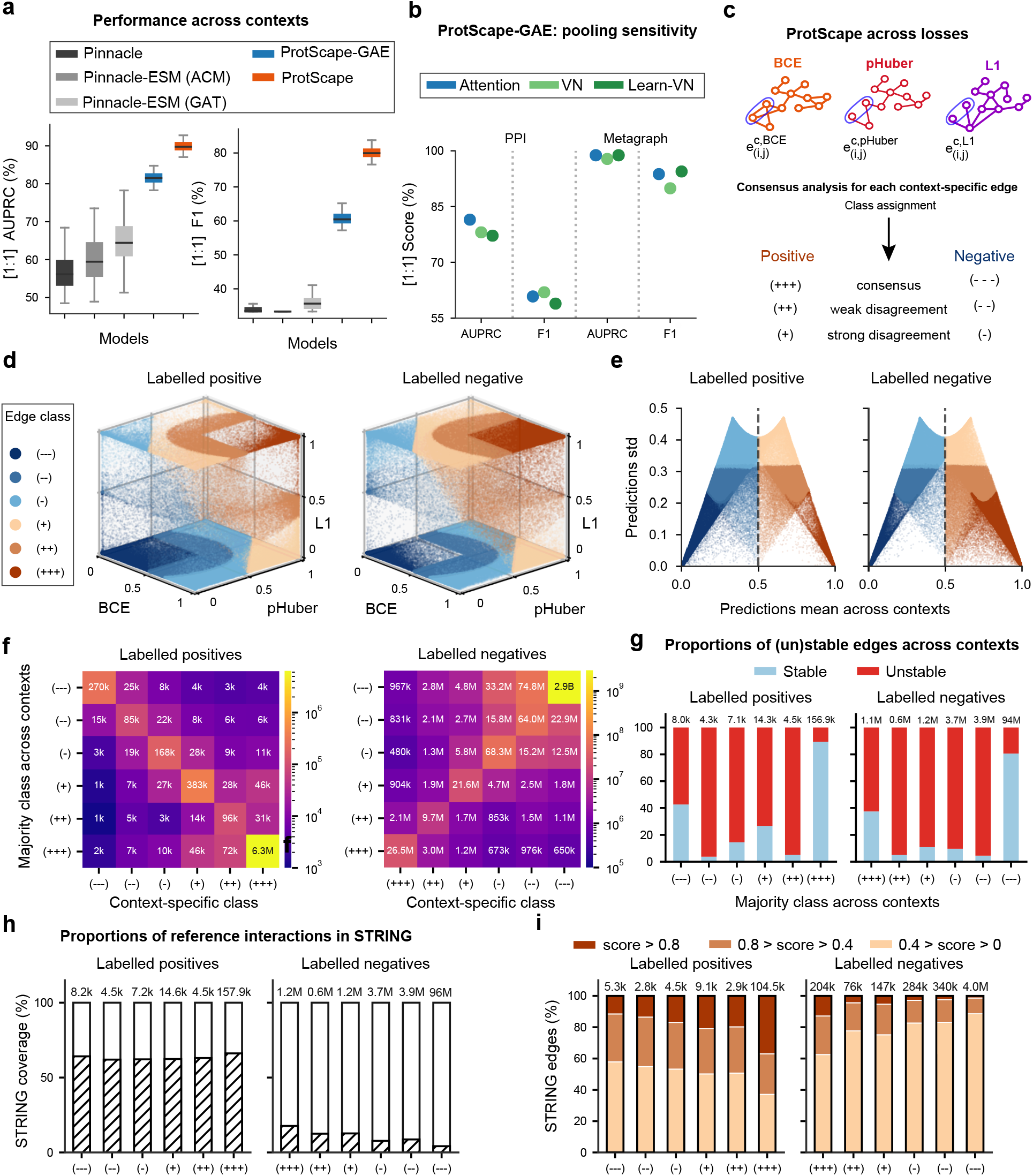
Extended evaluation of interaction reconstruction, loss sensitivity and STRING support. **a**, Distribution of held-out Cell-PPI prediction performance across cellular contexts under balanced 1:1 evaluation, comparing Pinnacle variants, ProtScape-GAE and ProtScape. Performance is reported as AUPRC and macro-F1. **b**, Sensitivity of ProtScape-GAE to protein-to-context pooling, comparing attention pooling, a zero-initialised virtual node (VN) and a learned context-specific VN for PPI and metagraph reconstruction. **c**, Consensus classes derived from ProtScape predictions obtained with BCE, pHuber and L1 reconstruction objectives. Each context-specific interaction is assigned to one of six classes ranging from consensus negative (—) to consensus positive (+++), with intermediate classes reflecting increasing disagreement across objectives. **d**, Joint distributions of BCE, pHuber and L1 prediction scores for labelled positive and negative interaction–context pairs, coloured by consensus class. **e**, Mean and variability of interaction predictions across cellular contexts for labelled positive and negative protein pairs, stratified by consensus class. Dashed lines indicate a mean prediction score of 0.5. **f**, Relationship between majority consensus class across contexts and context-specific class assignments. Heatmaps show the number of context-specific occurrences in each class for labelled positive and negative interactions. **g**, Cross-context stability of interactions according to majority consensus class. Bars show the proportions retaining the same class across all contexts or changing class between contexts; numbers indicate unique interactions per majority class. **h**, STRING coverage of interactions according to majority consensus class, shown separately for labelled positive and negative protein pairs. Hatched areas denote interactions represented in STRING; numbers indicate interactions per class. **i**, Distribution of STRING combined scores among STRING-supported interactions in each majority consensus class, grouped into low (0–0.4), intermediate (0.4–0.8) and high (*>*0.8) evidence categories. Numbers indicate STRING-supported interactions per class.

**Extended Data Fig. 3.**
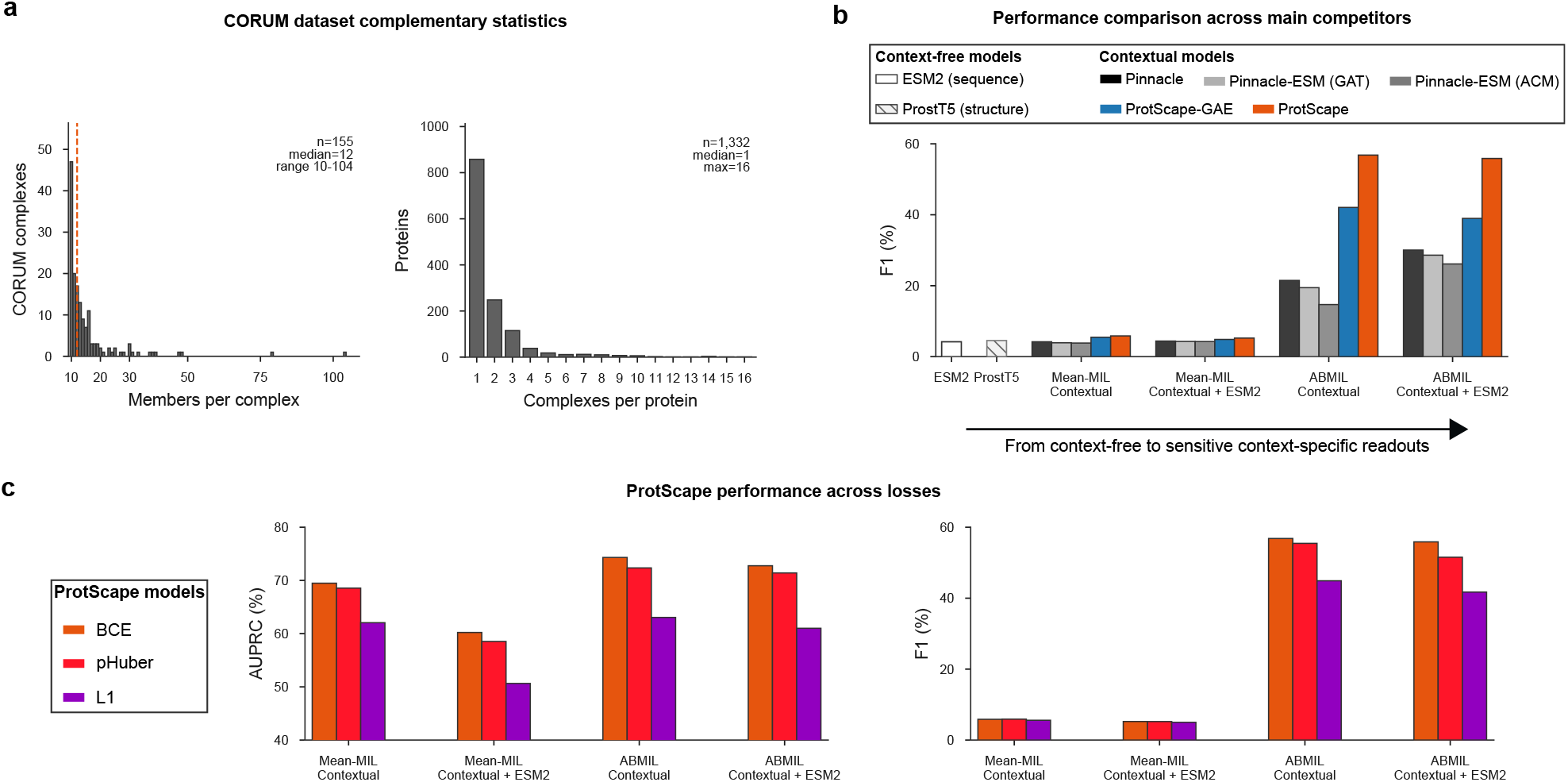
Extended evaluation of context aggregation and pretraining objectives on the CORUM protein-complex benchmark. **a**, CORUM benchmark composition after processing. Left, distribution of proteins per retained complex; right, distribution of retained complex memberships per protein. Dashed line indicates the median complex size. **b**, Protein-complex prediction performance for context-free ESM2 and ProstT5 representations and contextual Pinnacle, ProtScape-GAE and ProtScape representations using mean multiple-instance learning (Mean-MIL) or attention-based multiple-instance learning (ABMIL), with or without ESM2 late fusion. Bars report test F1. **c**, Sensitivity of ProtScape protein-complex prediction to the PPI reconstruction objective used during pretraining. Test AUPRC (left) and F1 (right) are shown for BCE-, pHuber- and L1-pretrained models across Mean-MIL and ABMIL readouts, with or without ESM2 late fusion.

**Extended Data Fig. 4.**
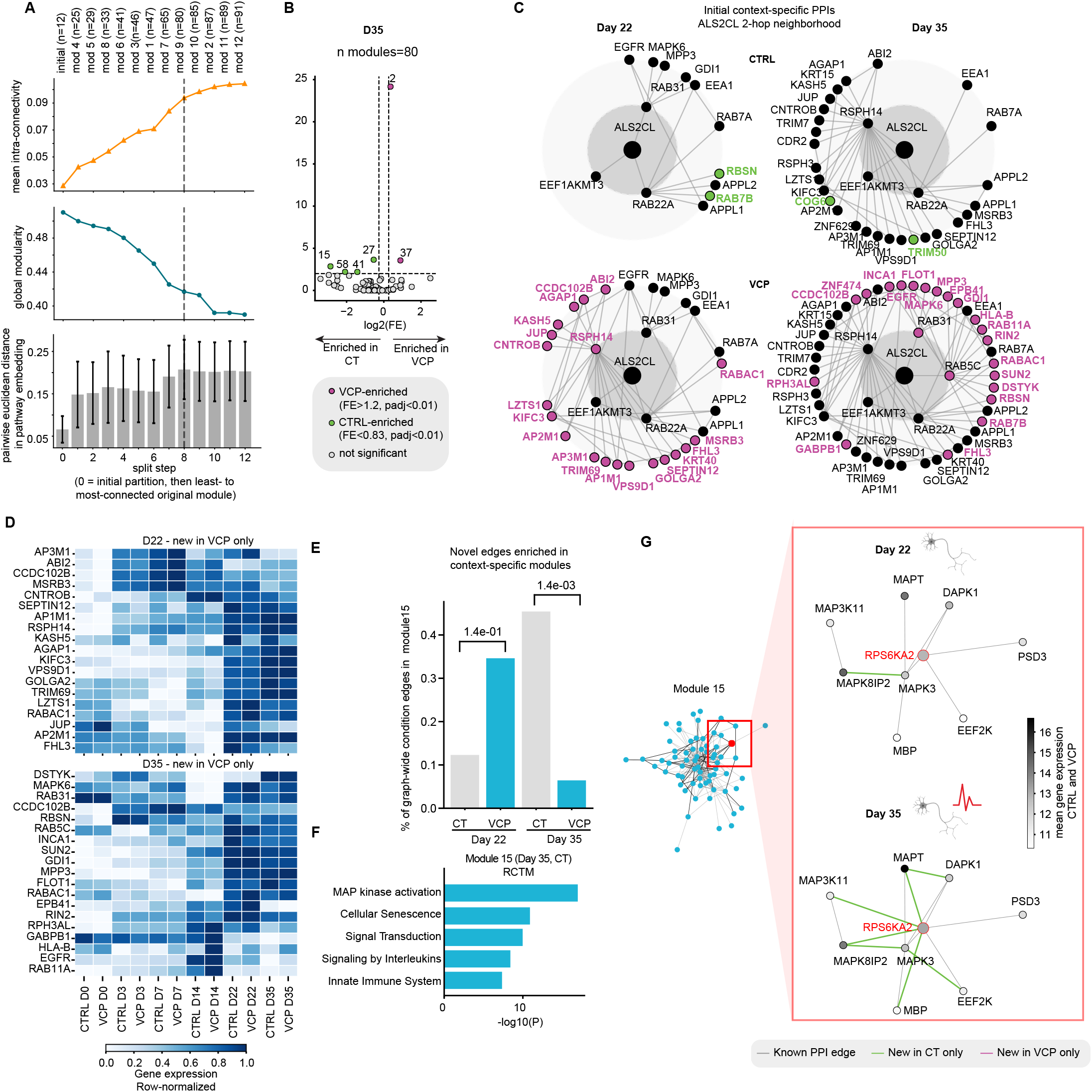
Extended analysis of condition-specific network rewiring during VCP-mutant motor-neuron differentiation. **a**, Robustness of iterative network partitioning, assessed by mean intra-module connectivity, global modularity and pairwise distance in pathway-embedding space across successive module splits. The selected partition of 80 modules is indicated by the dashed line. **b**, Enrichment of CTRL- and VCP-specific newly inferred interactions across modules at day 35. Fold enrichment compares their representation within each module with their graph-wide abundance; significantly enriched modules are highlighted (*P*_adj_ *<* 0.01; fold enrichment *>* 1.2 for VCP or *<* 0.83 for CTRL). **c**, Initial Cell-PPI topology surrounding ALS2CL in CTRL and VCP-mutant cells at days 22 and 35. Networks show the two-hop ALS2CL neighbourhood before ProtScape training, with condition-associated differences highlighted. **d**, Row-normalised expression profiles of proteins involved in VCP-specific newly inferred interactions at days 22 and 35 across motor-neuron differentiation. **e**, Fraction of graph-wide condition-specific interactions assigned to module 15 at days 22 and 35. *P* values compare the relative representation of CTRL- and VCP-specific interactions in the module using FDR-corrected Fisher’s exact tests. **f**, Reactome enrichment of module 15, highlighting MAP kinase activation and related signalling pathways. **g**, RPS6KA2-centred interaction neighbourhood within module 15 at days 22 and 35. Grey edges denote reference interactions and green edges interactions selectively inferred in CTRL cells; node shading indicates mean gene expression across CTRL and VCP conditions.

**Extended Data Fig. 5.**
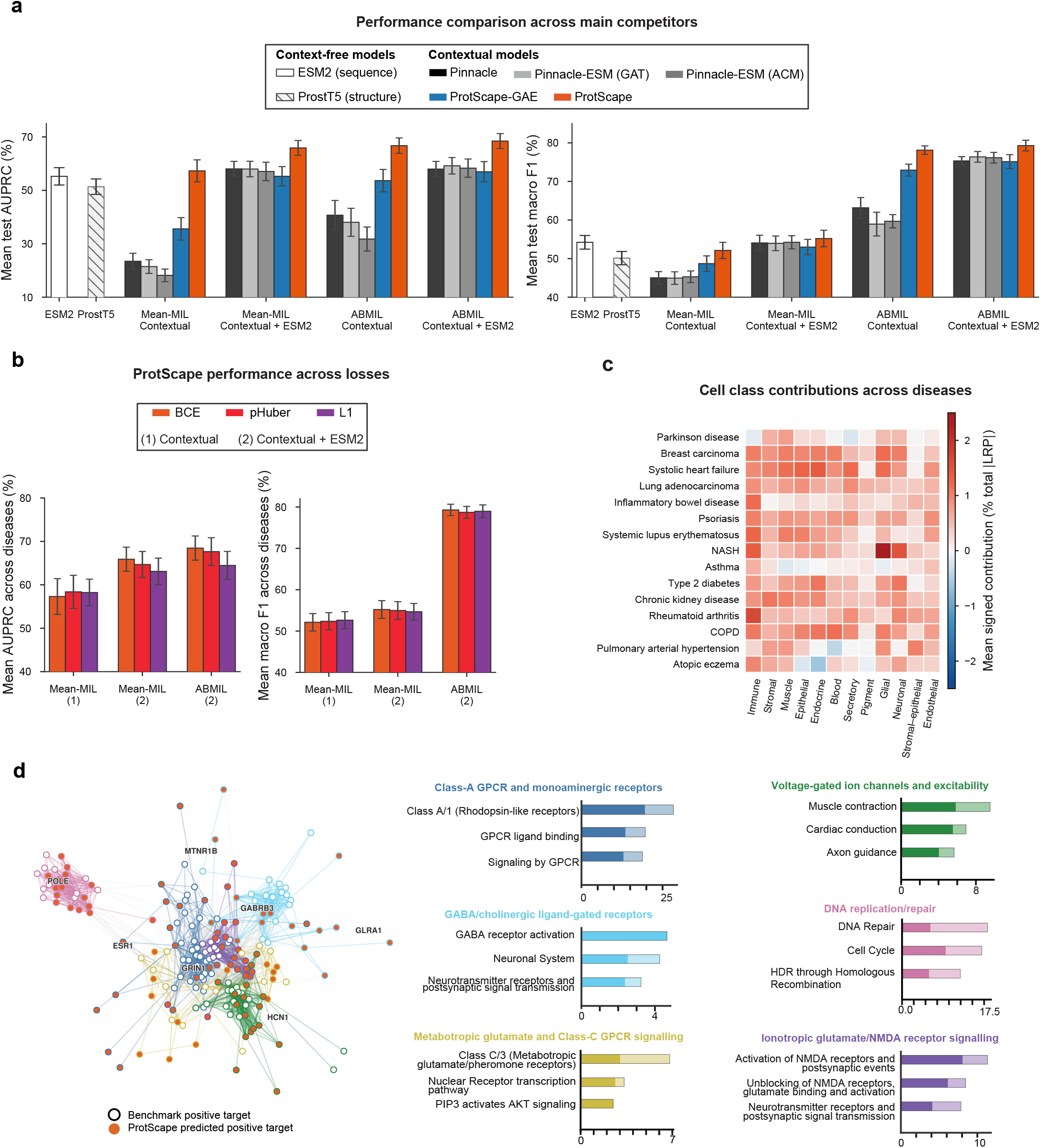
Complementary analyses of therapeutic-target prioritisation. **a**, Therapeutic-target prediction performance for context-free protein foundation models and contextual representations using mean multiple-instance learning (Mean-MIL) or attention-based multiple-instance learning (ABMIL), with or without ESM2 late fusion. Bars show mean test macro-F1 across 15 diseases; error bars denote s.e.m. **b**, Sensitivity of therapeutic-target prediction to the interaction-reconstruction objective used during ProtScape pretraining. Mean test AUPRC (left) and macro-F1 (right) are shown for models pretrained with binary cross-entropy (BCE), partially Huberized BCE (pHuber) or L1 reconstruction losses across contextual readouts; error bars denote s.e.m. across diseases. **c**, Disease- and cell-class-level interpretation of held-out therapeutic-target predictions recovered by ProtScape. Heatmap shows mean signed layer-wise relevance propagation (LRP) contributions, expressed as a percentage of total absolute relevance; red and blue indicate positive and negative contributions, respectively. **d**, STRING experimental-interaction network linking benchmark-positive Parkinson’s disease targets and ProtScape-nominated candidates, partitioned into six weighted Leiden communities. Adjacent bars show representative Reactome terms enriched within each community relative to the plotted 200-node network; bar length indicates −log_10_(FDR), with dark and light segments denoting benchmark-positive targets and predicted candidates, respectively.

**Extended Data Table. 1.**
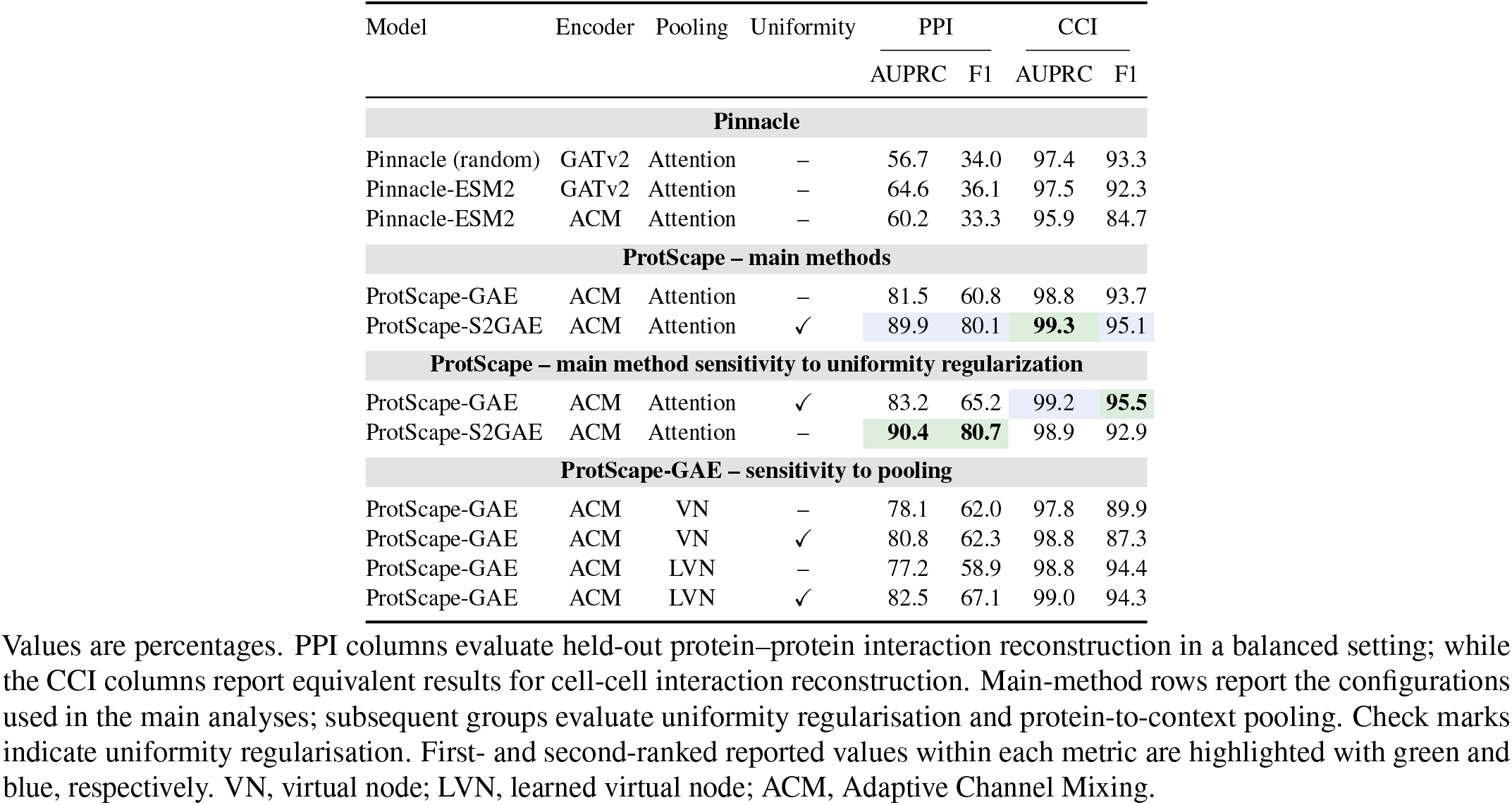
Pretraining performance and architectural ablations.

**Extended Data Table. 2.** CORUM complex-membership prediction across model readouts and pretraining ablations.

| Model | Encoder | Pooling | Uniformity | Context-free embeddings | Contextual instances |  |  | Contextual instances + ESM2 late fusion |  |  |
| --- | --- | --- | --- | --- | --- | --- | --- | --- | --- | --- |
|  |  |  |  | Linear probe | Mean-pool LR | ABMIL | ABMIL–PDL | Mean-pool LR | ABMIL | ABMIL–PDL |
| Sequence baselines |  |  |  |  |  |  |  |  |  |  |
| ESM2 | – | – | – | 34.4 (1.7) | – | – | – | – | – | – |
| ProstT5 | – | – | – | 34.5 (2.4) | – | – | – | – | – | – |
| Pinnacle |  |  |  |  |  |  |  |  |  |  |
| Pinnacle (random) | GATv2 | Attention | – | – | 17.2 (0.8) | 30.0 (1.5) | **44.9 (2.2)** | 36.6 (1.5) | 36.6 (2.7) | 43.1 (2.0) |
| Pinnacle-ESM2 | GATv2 | Attention | – | – | 15.1 (1.4) | 23.7 (2.7) | 38.2 (1.4) | 36.2 (1.5) | 36.9 (0.4) | **39.2 (0.9)** |
| Pinnacle-ESM2 | ACM | Attention | – | – | 11.3 (0.4) | 22.0 (0.7) | 28.8 (1.4) | 34.4 (1.7) | 35.0 (1.6) | **35.9 (1.1)** |
| ProtScape – main methods |  |  |  |  |  |  |  |  |  |  |
| ProtScape-GAE | ACM | Attention | – | – | 48.5 (3.0) | 51.3 (1.1) | **57.5 (1.1)** | 45.5 (0.6) | 50.6 (1.8) | 55.1 (1.4) |
| ProtScape-S2GAE | ACM | Attention | ✓ | – | 69.5 (1.4) | 70.4 (1.8) | **74.3 (0.7)** | 60.2 (1.1) | 69.1 (1.3) | **72.8 (0.8)** |
| ProtScape – main method sensitivity to uniformity regularization |  |  |  |  |  |  |  |  |  |  |
| ProtScape-GAE | ACM | Attention | ✓ | – | – | – | – | – | – | **62.9 (1.1)** |
| ProtScape-S2GAE | ACM | Attention | – | – | – | – | – | – | – | **71.5 (1.5)** |
| ProtScape-GAE – sensitivity to pooling |  |  |  |  |  |  |  |  |  |  |
| ProtScape-GAE | ACM | VN | – | – | – | – | – | – | – | **62.0 (2.1)** |
| ProtScape-GAE | ACM | VN | ✓ | – | – | – | – | – | – | **56.6 (1.3)** |
| ProtScape-GAE | ACM | LVN | – | – | – | – | – | – | – | **56.4 (1.0)** |
| ProtScape-GAE | ACM | LVN | ✓ | – | – | – | – | – | – | **58.0 (0.8)** |
| ProtScape-S2GAE – sensitivity to pretraining loss |  |  |  |  |  |  |  |  |  |  |
| ProtScape-S2GAE (pHuber) | ACM | Attention | ✓ | – | 68.6 (1.1) | 68.7 (1.4) | **72.4 (1.6)** | 58.5 (0.7) | 66.6 (1.2) | 71.4 (0.9) |
| ProtScape-S2GAE (L1) | ACM | Attention | ✓ | – | 62.1 (2.1) | 57.5 (2.8) | **63.0 (1.5)** | 50.6 (1.4) | 56.1 (2.2) | 61.0 (1.7) |

Values report test AUPRC (%), averaged across five fold-specific models and equally across complexes. Parentheses show sample SD across the five models. Sequence baselines use context-free embeddings; contextual readouts are evaluated with or without ESM2 late fusion. Main-method rows report the configurations used in the main analyses; subsequent groups evaluate uniformity regularization, protein-to-context pooling and alternative PPI reconstruction losses. Check marks indicate uniformity regularization; dashes denote unreported results. First- and second-ranked values across all models and readouts are highlighted with green and blue, respectively. VN, virtual node; LVN, learned virtual node; ACM, Adaptive Channel Mixing; LR, logistic regression; ABMIL, attention-based multiple-instance learning; PDL, progressive-dropout layers.

**Extended Data Table. 3.** Details on therapeutic target prediction datasets.

| Disease | Ontology ID | Abbrev. | Positives | Negatives |
| --- | --- | --- | --- | --- |
| Inflammatory bowel disease | EFO_0003767 | IBD | 126 | 1360 |
| Rheumatoid arthritis | EFO_0000685 | RA | 173 | 1327 |
| Breast carcinoma | EFO_0000305 | BrCa | 176 | 642 |
| Systolic heart failure | EFO_1001207 | SHF | 59 | 1928 |
| Lung adenocarcinoma | EFO_0000571 | LUAD | 52 | 1438 |
| Psoriasis | EFO_0000676 | Psor. | 170 | 1233 |
| Pulmonary arterial hypertension | EFO_0001361 | PAH | 63 | 1746 |
| Type 2 diabetes mellitus | MONDO_0005148 | T2D | 202 | 1364 |
| Systemic lupus erythematosus | MONDO_0007915 | SLE | 126 | 1623 |
| Atopic eczema | EFO_0000274 | Ecz. | 64 | 1696 |
| Asthma | MONDO_0004979 | Asthma | 112 | 1523 |
| Chronic obstructive pulmonary disease | EFO_0000341 | COPD | 154 | 1655 |
| Non-alcoholic steatohepatitis | EFO_1001249 | NASH | 62 | 1884 |
| Chronic kidney disease | EFO_0003884 | CKD | 190 | 1466 |
| Parkinson disease | MONDO_0005180 | PD | 107 | 1372 |

Ontology identifiers specify the root disease terms used to aggregate Open Targets evidence. Counts include global-interactome proteins with both ProtScape and ESM2 embeddings. Positives have phase 3 or later evidence, or completed phase 2 evidence; negatives are druggable proteins without comparable indication-specific evidence.

**Extended Data Table. 4.**
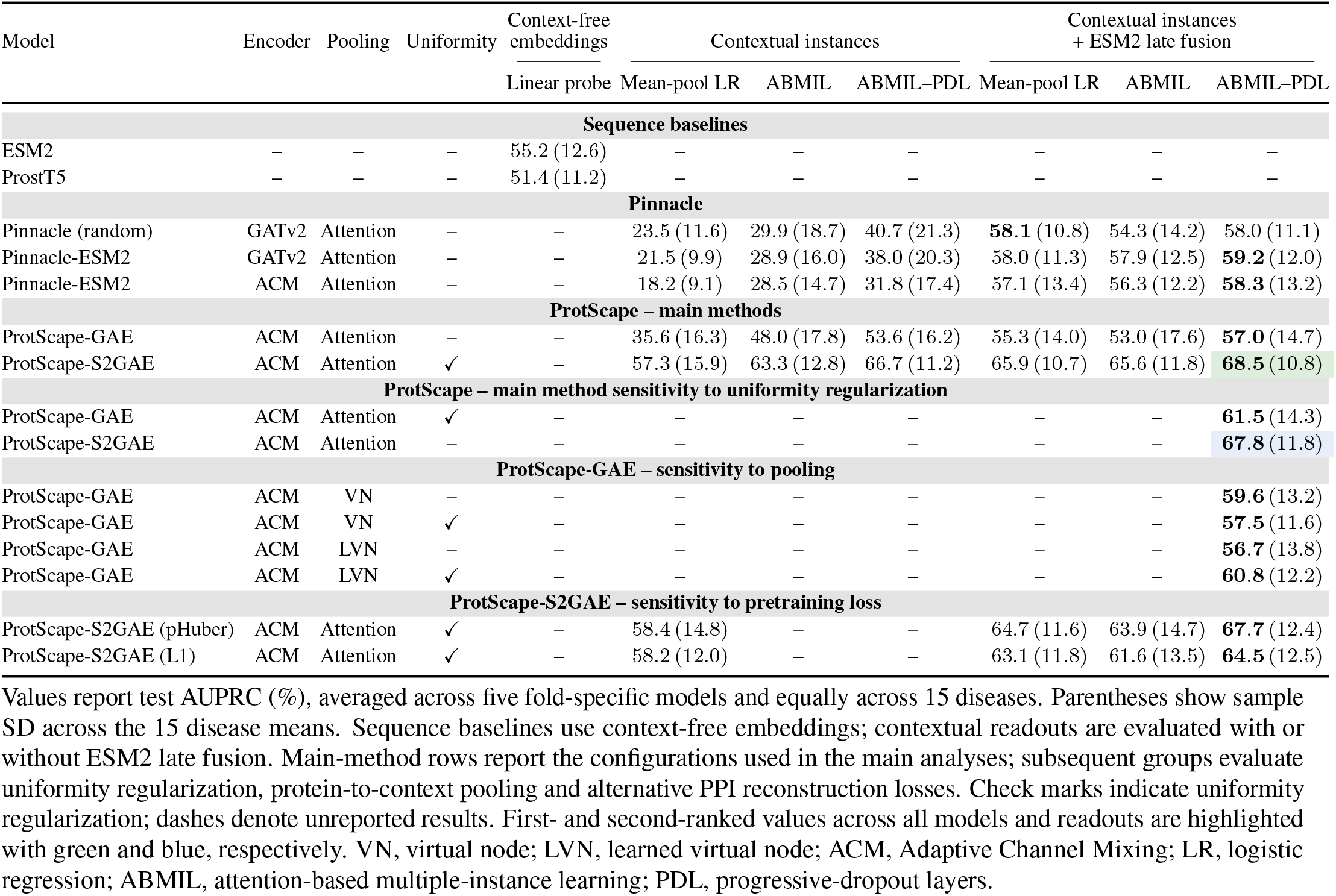
Therapeutic-target prediction across model readouts and pretraining ablations.

**Extended Data Table. 5.** Disease-specific therapeutic-target prediction performance.

| Disease | ESM2 | ProstT5 | Pinnacle<br>(random) | Pinnacle-ESM2<br>GATv2 | Pinnacle-ESM2<br>ACM | ProtScape<br>GAE | ProtScape<br>BCE | ProtScape<br>pHuber | ProtScape<br>L1 |
| --- | --- | --- | --- | --- | --- | --- | --- | --- | --- |
| **Mean** | 55.2 (12.6) | 51.4 (11.2) | 58.0 (11.1) | 59.2 (12.0) | 58.3 (13.2) | 57.0 (14.7) | **68.5 (10.8)** | 67.7 (12.4) | 64.5 (12.5) |
| IBD | 65.3 (5.5) | 55.2 (6.6) | 65.0 (7.1) | 67.2 (6.3) | 66.2 (5.9) | 52.8 (9.6) | **71.6 (5.6)** | 69.9 (4.9) | 64.2 (6.8) |
| RA | 54.5 (6.0) | 43.9 (1.4) | 53.5 (5.3) | 54.7 (8.4) | 56.7 (5.9) | 46.5 (6.7) | **62.6 (3.7)** | 61.3 (4.6) | 58.2 (6.9) |
| Breast cancer | 76.7 (4.3) | 68.7 (5.1) | 79.1 (6.9) | 82.1 (3.3) | 79.6 (2.8) | 82.1 (2.3) | 83.6 (4.4) | 83.1 (3.8) | **89.0 (0.6)** |
| Systolic HF | 50.4 (5.2) | 56.2 (2.8) | 61.2 (6.4) | 66.7 (7.1) | 63.0 (4.0) | 71.8 (10.2) | 80.1 (1.1) | **80.5 (1.2)** | 70.8 (1.7) |
| LUAD | 54.4 (7.3) | 38.6 (5.2) | 59.5 (4.2) | 48.5 (20.4) | 59.6 (6.0) | 58.6 (10.0) | **76.3 (3.8)** | 74.0 (3.2) | 59.7 (6.4) |
| Psoriasis | 52.2 (6.2) | 61.4 (6.0) | 55.0 (4.6) | 55.0 (5.4) | 54.9 (5.5) | 60.2 (6.1) | 70.7 (1.9) | **71.4 (2.0)** | 67.5 (3.6) |
| PAH | 35.5 (4.6) | 34.6 (5.9) | 49.2 (7.2) | 50.5 (8.5) | 44.0 (11.6) | 25.0 (3.9) | **54.4 (1.7)** | 52.6 (3.6) | 45.5 (4.0) |
| T2D | 55.6 (3.0) | 61.5 (2.1) | 57.6 (4.3) | 61.4 (4.2) | 57.4 (2.7) | 60.8 (4.7) | 63.5 (2.0) | **69.1 (1.6)** | 68.4 (2.5) |
| SLE | 56.0 (3.6) | 55.5 (6.4) | 64.0 (4.7) | 62.9 (4.9) | 56.7 (6.8) | 59.5 (4.1) | **68.6 (3.5)** | 65.7 (3.1) | 62.8 (2.7) |
| Atopic eczema | 34.1 (4.9) | 35.6 (4.2) | 39.7 (7.2) | 36.2 (4.0) | 32.6 (3.9) | 37.6 (6.0) | **50.7 (6.2)** | 40.1 (4.1) | 46.9 (4.1) |
| Asthma | 61.1 (1.6) | 49.8 (3.2) | 51.1 (4.6) | 64.8 (5.8) | 66.9 (6.9) | 54.2 (5.0) | 64.3 (7.4) | **72.7 (6.0)** | 66.2 (8.1) |
| COPD | 40.3 (4.3) | 40.2 (1.7) | 40.9 (3.7) | 47.1 (5.9) | 41.5 (3.4) | 46.8 (3.6) | **60.4 (2.1)** | 60.4 (2.8) | 51.9 (4.4) |
| NASH | 69.8 (4.5) | 61.7 (3.5) | 60.6 (21.6) | 62.1 (19.3) | 64.2 (19.8) | 60.3 (9.5) | 66.0 (9.0) | 62.1 (14.3) | **72.9 (10.3)** |
| CKD | 50.3 (2.1) | 43.4 (3.8) | 55.7 (6.5) | 51.5 (4.8) | 49.7 (4.0) | 59.0 (2.5) | **63.3 (3.5)** | 61.5 (3.4) | 56.3 (3.4) |
| Parkinson disease | 72.4 (4.3) | 64.1 (9.1) | 77.5 (8.2) | 77.4 (4.9) | 81.9 (3.4) | 79.4 (2.4) | **90.9 (3.3)** | 90.4 (3.3) | 86.5 (5.0) |

Values report test AUPRC (%) averaged across five fold-specific models on the corresponding fixed test set. Mean is the average across 15 diseases. Parentheses show sample SD across five models for disease rows and across 15 disease means for the Mean row. ESM2 and ProstT5 use sequence-only linear probes; contextual models use ABMIL–PDL with ESM2 late fusion. Settings were selected by validation AUPRC. Green highlights the highest AUPRC in each row. ABMIL, attention-based multiple-instance learning; PDL, progressive-dropout layers. IBD, inflammatory bowel disease; RA, rheumatoid arthritis; HF, heart failure; LUAD, lung adenocarcinoma; PAH, pulmonary arterial hypertension; T2D, type 2 diabetes mellitus; SLE, systemic lupus erythematosus; COPD, chronic obstructive pulmonary disease; NASH, non-alcoholic steatohepatitis; CKD, chronic kidney disease.

